# Stress Granule Sequestration of CCR4–NOT Promotes Poly(A) Lengthening of Stress-Survival Transcripts

**DOI:** 10.1101/2025.02.04.636058

**Authors:** Yi-Cheng Tsai, Hiroyuki Uechi, Sean R. Millar, Eileigh Kadijk, Alon Minkovich, John Paul Tsu Ouyang, Karl J. Schreiber, Jie Qi Huang, Carina A. Lyons, Vesal Kasmaeifar, Zhixin Zhang, Rico Barsacchi, Claudia Möbius, Antje Janosch, Jonathan C. Savage, Olivia S. Rissland, Hannes Röst, Anthony A. Hyman, Ji-Young Youn

**Author notes:** Senior Author (Anthony A. Hyman, Ji-Young Youn). Graduate School of Pharmaceutical Sciences, Kyoto University, 606-8501 Kyoto, Japan (Hiroyuki Uechi). Data deposition: Mass spectrometry data have been deposited in the Mass Spectrometry Interactive Virtual Environment (MassIVE, http://massive.ucsd.edu). MassIVE ID: MSV000095953 (ftp://massive.ucsd.edu/v08/MSV000095953/) MassIVE ID: MSV000095944 (ftp://massive.ucsd.edu/v08/MSV000095944/) MassIVE ID: MSV000098662 (ftp://massive-ftp.ucsd.edu/v10/MSV000098662/) MassIVE ID: MSV000100193 (ftp://massive-ftp.ucsd.edu/v11/MSV000100193/).

## Abstract

Cells adapt to stress by rewiring their post-transcriptional gene regulation. Stress granules—biomolecular condensates composed of polyadenylated RNAs and RNA-binding proteins—are implicated in this process, yet their precise functional roles remain debated. To address this, we mapped the dynamic proteomic landscapes of stress granules formed under oxidative and hyperosmotic stress using multi-bait BioID proximity profiling coupled with quantitative mass spectrometry. This analysis revealed context-specific remodeling of proximal interaction networks and identified a conserved, stress-dependent shift in association with the CCR4–NOT deadenylase complex. A complementary genome-wide chemical genetic screen further implicated CCR4-NOT in stress granule biology, showing that reduced CCR4–NOT activity bypassed lipoamide-mediated inhibition of stress granule assembly. Microscopy showed sequestration of the CCR4-NOT complex into stress granules, and global transcriptomic analyses revealed that this relocalization promotes poly(A) tail lengthening and increased abundance of stress-induced survival transcripts. Together, integration of proteomics, chemical genetics, and transcriptomics uncovers a spatial mechanism by which stress granule assembly promotes cellular adaptation to stress through sequestration of CCR4-NOT from the cytosol and consequent remodeling of post-transcriptional regulation.

## MAIN

Living organisms have evolved mechanisms to survive unfavourable conditions. At the cellular level, adaptation to stress involves rewiring transcriptional programming, translation control, signaling, metabolism, and the reorganization of subcellular proteomes. Cells shift biological processes to conserve resources during stress, slowing global transcription and translation to minimize energy consumption while actively transcribing genes involved in stress responses^1,2^.

Cellular adaptation involves changes in the subcellular proteome organization^3-5^. Advances in spatial proteomics, including imaging-based microscopy and mass spectrometry, have enabled the identification of proteins localized to specific subcellular compartments^4,6-12^. During stress, proteins dynamically exchange between compartments^5,13-16^; however, assigning functional roles to these localization changes remains challenging.

The re-organization of subcellular proteomes often involves the formation of membraneless organelles or biomolecular condensates, which concentrate biomolecules through liquid-liquid phase separation^17-20^. By enriching specific biomolecules, biomolecular condensates can increase or decrease biochemical reactions, buffer protein concentrations, sense environmental changes, and provide mechanical forces^21-25^. Stress granules are cytosolic biomolecular condensates or RNA granules that transiently form in response to various stress conditions, often as a product of global translation shutdown and release of mRNAs from the polysome pool^26,27^. Stress granules host hundreds of proteins and several thousand transcripts^28-31^ and are central to stress response, linking translation control to signaling and immune responses^29,32,33^. They host antiviral responses during viral infection^27^ and sequester regulators of cell death^34,35^ or cell growth pathway components during stress^36^.

Elucidating stress granule function is confounded by the variability observed depending on the cellular stress or cell type^29,37^. For example, depending on how translation is blocked, stress granules vary in their composition of translation initiation factors. Activation of the integrated stress response and eIF2α phosphorylation induces stress granules lacking eIF2-eIF5-Met-tRNA^Met^ ternary complex, while eIF4A poisons (e.g., Pateamine A) induce stress granules containing this ternary complex. Stress granules induced by chemotherapeutic drugs, such as selenite, UV and other xenobiotic agents, lack eIF3^29,38^. Lastly, hyperosmotic stress induces phospho-eIF2α-independent stress granules through molecular crowding; however, stress granules induced by hyperosmotic stress is poorly characterized. This underscores the importance of studying stress granules induced by distinct mechanisms.

A key aspect of the stress response is the upregulation of proteins required for adaptation, including chaperones and transcription factors that promote cell survival^26,39-43^. However, how the corresponding mRNAs evade mRNA decay pathways to maintain high abundance and preferential translation remains unclear.

As part of post-transcriptional regulation, modulation of poly(A) tail lengths is crucial for protecting mRNA from decay^44^. Poly(A) tail lengths are controlled by a balance between the action of cytoplasmic poly(A) RNA polymerase (*e.g.,* GLD2 and TENT5B) that elongate these tails and deadenylases (*i.e.,* CCR4-NOT, PAN2-3 complexes and PARN) that shorten them^44,45^. Among these three deadenylases in vertebrates ^44^, the CCR4-NOT complex is the principal deadenylation responsible for degrading poly(A)s^44-46^. According to a biphasic model of deadenylation, the PAN2-PAN3 complex initiates the poly(A) tail removal, while the CCR4-NOT complex trims the remaining tail close to the 3’-UTR^47^. The CCR4-NOT complex consists of eleven subunits, with a conserved core formed by two major modules: the NOT module (comprising the scaffolding protein CNOT1 and a CNOT2-CNOT3 heterodimer) and the catalytic module (one of the deadenylase CCR4 units, CNOT6 or CNOT6L, paired with one of the exonuclease CAF1 units, CNOT7 or CNOT8). The CNOT6, CNOT6L, CNOT7 and CNOT8 subunits perform the deadenylase activity^44^, while the NOT module (CNOT1, CNOT2, CNOT3) is critical for maintaining complex integrity, regulating the catalytic module’s activity, and recruiting the complex to specific mRNAs via specific interactions with RNA-binding or GW182 proteins^48-55^.

Acute stress induces poly(A) tail lengthening^56-59^. While stress-induced stabilization of transcripts has been linked to inhibiting deadenylation^58,60^, the molecular mechanisms behind stress-induced poly(A) tail lengthening remain elusive. Recent work has demonstrated that poly(A) tail extension is a prerequisite for stress granule assembly^61^, hinting at a potential link between stress response and the regulation of enzymes controlling poly(A) lengths. However, clear evidence connecting poly(A) tail regulatory pathways to stress responses remains limited.

Recently, multi-bait proteomics has become a powerful approach for studying protein localization within condensates^62,63^. In this study, we use multi-bait BioID time course experiments to map the proximal interactome of stress granule proteins, providing a resource that captures proteomic landscape changes during stress. Leveraging this resource, we demonstrate that interactions with the CCR4-NOT complex shift dynamically in response to stress. A complementary genome-wide CRISPR screen against stress granule-inhibiting chemical lipoamide revealed that the removal of CCR4-NOT stabilizes stress granules through poly(A) tail lengthening. Microscopy analysis of CCR4-NOT showed poor localization to P-bodies, contrary to its annotation, but strong localization to stress granules during stress. We propose that stress granules promote cellular adaptation by sequestering the CCR4–NOT complex, attenuating its deadenylase activity and enabling poly(A) tail lengthening that sustains transcripts important for cell survival.

## RESULTS

### Dynamic Changes in Proximal Interactions of Stress Granule Proteins

We examined the proximal interaction networks of stress granule proteins G3BP1 and FMR1 during oxidative stress. Previously, we reported that the proximal interactions of G3BP1 and EIF4A1 remained largely unchanged before and after stress granule assembly when using the slow biotin ligase BirA* ^63^. However, this approach involved a labeling period significantly longer than the known kinetics of stress granule assembly. To capture these interactions more accurately, we employed the faster biotin ligase, miniTurbo^64^, which enables proximal labeling within 15-30 min, compared to the 3–24 h required by BirA*. We integrated expression constructs of G3BP1 and FMR1, each fused to miniTurbo-3xFLAG (miniT), into the inducible Flp-In locus T-REx HEK293 system. The expression levels of miniT-tagged proteins were comparable to their endogenous counterparts (Figure S1A), increasing total protein abundance by about two-fold. Since G3BP1 is known to scaffold stress granule assembly, we checked if miniT-G3BP1 induction induces stress granule assembly in the absence of stress and found no visible stress granules (Figure S1B).

To assess proximal interaction changes upon stress granule assembly, we induced bait protein expression in biotin-depleted media and pulse-labeled for 30 min in either no stress condition (T1) or 5 min into oxidative stress induced with 0.5mM sodium arsenite (III) (T2; Figure 1A; see Methods). Proteins biotinylated during T1 displayed a general cytoplasmic distribution, whereas during T2, they were highly enriched in cytosolic puncta, colocalizing with the stress granule marker TIAR (Figure S1B).

**Figure 1.**
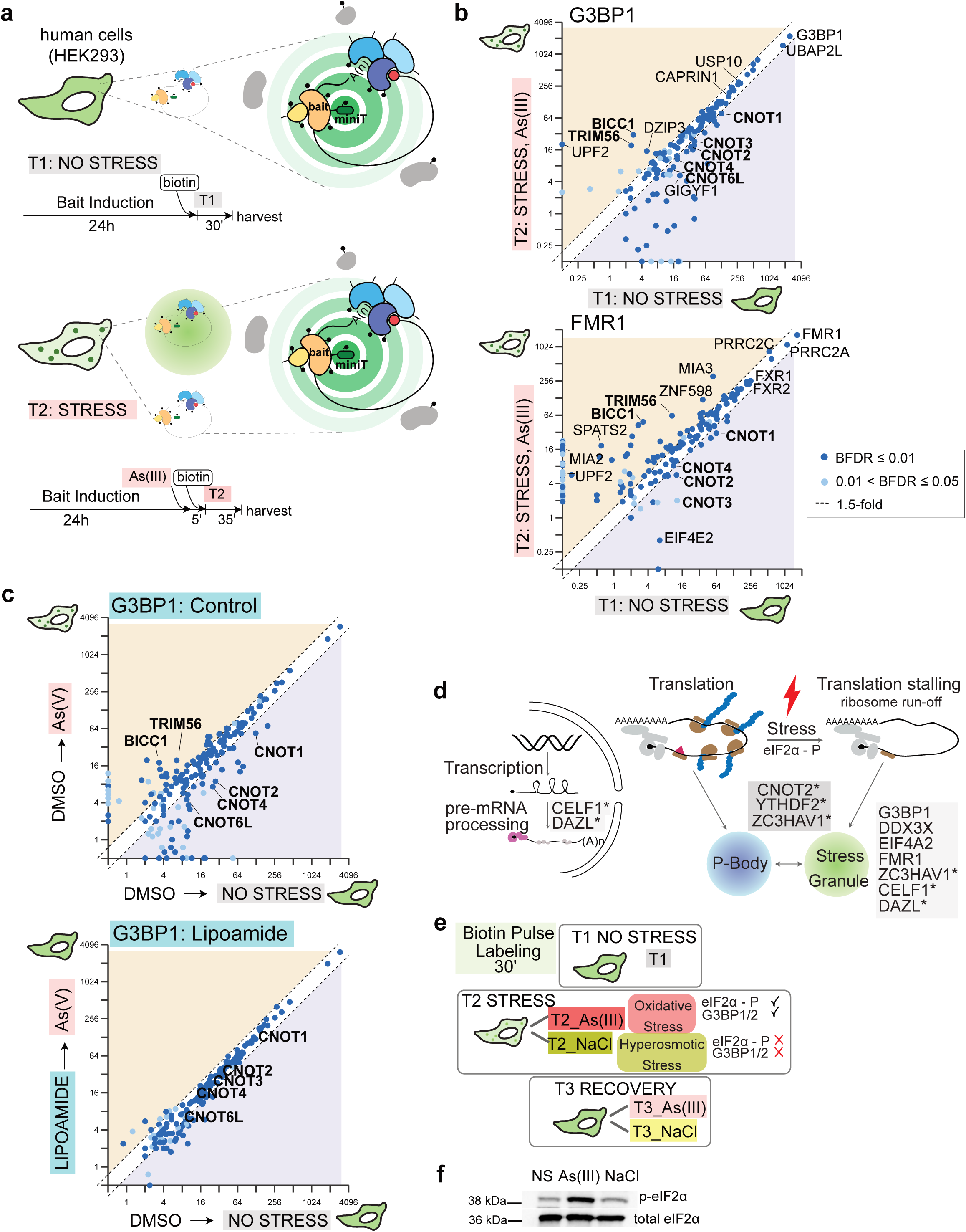
BioID of Stress Granules during steady state and stress. (**a**) Schematics show experimental workflow of stress granule proximity labelling no stress (top) or stressed state with oxidative stress (bottom) in human cells. (**b**) Quantitative comparison of average spectra (log_2_ transformed) of high-confidence interactors identified by G3BP1 (top) and FMR1 (bottom) during no stress *vs*. stress state (0.5mM sodium arsenite (III)) using ProHits-Viz^131^. (**c**) Comparative analyses of carrier pre-treated stress BioID (top) *vs.* lipoamide pre-treated stress BioID (bottom). The condition-condition plot shows the abundance of high-confidence interactors identified by G3BP1 in each condition. (**d**) Schematics of nine high-information baits and their subcellular location. Baits with asterisks localize to multiple compartments. (**e**) Diagram of stress BioID experimental workflow. Proximity labelling is performed during no stress, stress (arsenite, As (III); oxidative stress, or NaCl; hyperosmotic stress), or recovery periods for 30 min. (**f**) Western blot shows eIF2α phosphorylation status in cells used for BioID (HEK293 Flp-In T-REx cells) during no stress (NS), oxidative (As(III)) or hyperosmotic stress (NaCl) used for BioID.

Biological duplicates of G3BP1, FMR1, and negative controls (miniT alone, miniT-GFP, and parental cells) were processed for BioID coupled to mass spectrometry (MS) followed by SAINTexpress analysis^65,66^ to determine high-confidence proximal interactors (BFDR ≤0.05). G3BP1 and FMR1 identified 137 and 102 high-confidence interactors during no stress and 107 and 139 during stress, respectively (Table S1, BFDR ≤0.05).

Most proximal interactors remained similar in abundance as reported^37,63^, confirming pre-existing proximal interactions between stress granule components in the absence of microscopically visible condensates. However, the improved labeling kinetics of the miniTurbo enzyme revealed a higher fraction of interactors displaying stress-dependent abundance changes. Approximately 4% and 30% of G3BP1 interactors, as well as 34% and 15% of FMR1 interactors, displayed ≥1.5-fold increase or decrease in abundance, respectively (Figure 1B). Common changes in G3BP1 and FMR1’s stress BioID profiles included shared increased interactors (*e.g.,* BICC1, TRIM56) and decreased interactors (*e.g.,* CNOT1, CNOT2, CNOT3, CNOT4). These findings indicate coordinated dynamic rearrangements in proximal interaction networks as G3BP1 and FMR1 condense to form stress granules.

### Stress-Induced Proximal Interaction Changes Are Reversed by Stress Granule Inhibitor, Lipoamide

Most stress-induced changes common to G3BP1 and FMR1 are abundance changes (∼1.5-3-fold changes in spectral counts) rather than their complete situational absence or presence (Figure 1B). To examine if these observed abundance changes reflect dynamic molecular interactions associated with stress granule assembly, we repeated the BioID experiment under the stress granule-inhibiting effects of lipoamide. Lipoamide is an antioxidant that prevents stress granule assembly during oxidative stress by blocking the oxidation of proteins. This mechanism primarily acts on intrinsically disordered proteins, such as SRSF1 and SFPQ. The redox state of the latter influences stress granule formation^67^.

Titration tests of potassium arsenate (As(V)), and lipoamide were used to identify effective concentrations for inhibiting stress granule assembly in HEK293 BioID miniT-GFP and miniT-G3BP1 cells. Note that all experiments involving lipoamide were used alongside the pentavalent form of As(V) (*e.g.,* potassium arsenate or sodium arsenate), which is a weaker oxidizer than the trivalent form of As(III) (*e.g.,* sodium arsenite). It was necessary to increase lipoamide concentration to fully inhibit stress granule assembly as potassium arsenate levels rose (Figure S1C). Based on these tests, cells were pre-treated with 100 μM lipoamide before inducing stress with 1 mM potassium arsenate (As(V)), followed by 30 min biotin labeling. Pre-treatment with a DMSO control followed by As(V) stress induced abundance changes in G3BP1 proximal interactions (*e.g.,* TRIM56, BICC1, CCR4-NOT complex components) that paralleled those seen during As(III) stress (Figure 1C, top). However, when pre-treated with lipoamide, interaction profiles remained at no stress levels and failed to recapitulate any stress-induced changes (Figure 1C, bottom). Further, lipoamide pre-treatment had no effect on G3BP1’s interaction profile during no stress (Figure S1D). We propose that lipoamide’s modulation of the redox state of stress granule proteins inhibits stress-induced changes in the G3BP1 proximal interaction network. These findings indicate that the observed abundance changes in G3BP1 and FMR1’s BioID profiles reflect dynamic molecular rearrangements during stress granule assembly.

### Mapping the Dynamic Stress Granule Interactome During Oxidative and Hyperosmotic Stress

The oxidative stress-induced changes in G3BP1’s interaction profile were similarly observed during FMR1 BioID. We adopted a multi-bait experimental design to extensively define the dynamic stress granule proximal interaction network. Nine baits known to localize to stress granules, or their related condensates, were selected with the criteria of maximizing the identification of stress granule proteins. Combining these baits (CELF1, DAZL, CNOT2, YTHDF2, ZC3HAV1, G3BP1, DDX3X, EIF4A2 and FMR1; Figure 1D; see Methods) was predicted to recover 60% of high-confidence interactors identified in no stress BioID dataset using 58 BirA*-conjugated baits^63^ (with the assumption that miniTurbioID provides similar labeling profiles), and 87% of the literature-curated stress granule proteome (Figure S1E, Table S2).

To investigate the effect of stress context in stress granules’ interaction networks, we incorporated two stressors - oxidative and hyperosmotic – with distinct stress granule induction mechanisms into our BioID screens. This approach aimed to identify both conserved and unique proteomic changes associated with distinct stress granule assembly pathways. Oxidative stress induces stress granules by the scaffolding of key nucleator proteins (G3BP1/2) and activation of the integrated stress response. Interfering with either of these, by means of knockout (G3BP1Δ, G3BP2Δ) or loss-of-function mutation (eIF2α S51A)^68,69^, is sufficient to curtail stress granule assembly. In contrast, hyperosmotic stress does not depend on G3BP1/2 nucleators or eIF2α phosphorylation^68-70^ (Figure 1E). Instead, the resulting cellular crowding during hyperosmotic stress drives stress granule nucleation^71,72^. In general, the cellular changes caused by oxidative *vs.* hyperosmotic stresses are distinct; oxidative stress creates reactive oxygen species, which damages cellular biomolecules and organelles^73-75^, while hyperosmotic stress induces immediate loss of water and activation of condensates inducing regulatory volume increase to compensate for volume loss^76,77^.

Accordingly, we performed separate BioID time course experiments to measure the dynamic rearrangement of the stress granule proximal interaction network during exposure and recovery from both stressors. We performed biotin labeling 5 min into oxidative stress for 30 min (T2_As(III)) and 1h after recovering from oxidative stress (T3_As(III)) based on our estimate of dissolution kinetics (Figure S2A) and published report^78^. For hyperosmotic stress, we noted small stress granules forming upon exposure to 0.2M NaCl, with growth observed up to 45 min into stress (Figure S2B). We observe robust eIF2α phosphorylation upon 30 min of exposure to oxidative stress and a moderate increase after 60 min of hyperosmotic stress (Figure 1F). Biotin labeling commenced 45 min into NaCl stress for 30 min (T2_NaCl), and we confirmed the enrichment of biotinylated proteins in NaCl-induced stress granules (Figure S2C). Upon removal of hyperosmotic stress, stress granules rapidly disassembled (Figure S2D). Consequently, we initiated biotin labeling immediately after hyperosmotic stress removal (T3_NaCl; Figure S2E). The differences in disassembly kinetics between oxidative (As(III)) and hyperosmotic (NaCl) stress correlated with faster translational resumption following hyperosmotic stress recovery (Figure S2F).

Nine baits localizing to stress granules or stress granules and P-bodies were stably and inducibly expressed in HEK293 cells in biotin-depleted media. The cells underwent 30 min of biotin pulse labelling at various time points (T1-3) under oxidative (As(III)) or hyperosmotic (NaCl) stress, along with negative controls (miniTurbo alone and miniTurbo-GFP) (Figure 1E, Figure S2E, S2G). We identified 5,605 high-confidence proximal interactions across 45 bait-stress conditions and 754 preys (BFDR ≤ 0.05 SAINTexpress, Table S3) using mass spectrometry data acquired in conventional <u>D</u>ata-<u>D</u>ependent <u>A</u>cquisition (DDA) mode, followed by SAINT analysis. Gene Ontology (GO) analysis revealed terms related to stress granules (cytoplasmic stress granule; GO:0010494, mRNA binding; GO:0003729, translation; GO:0006412), and unanticipated terms like microtubule organizing center (GO:0005815) and microtubule cytoskeleton (GO: 0015630) (Table S3). The significance of microtubules and motor proteins in stress granule assembly has been previously documented for stress granules induced by oxidative stress^79,80^. However, in this dataset, proximal interactions with microtubule proteins were encountered specifically during the T2_NaCl timepoint (example of DDX3X BioID in Figure S3A).

To see if these interactions with microtubule-associated proteins are unique to “mature” stress granules formed later in stress, we performed BioID in DDX3X-miniT at an earlier time point (5-35 min post-NaCl exposure) and found similar results. These interactions were reproduced during hyperosmotic stress induced by sorbitol (0.4M) (Figure S3B, Table S3), indicating that DDX3X’s increased association with microtubule and centrosome proteins is a general response to hyperosmotic stress. To assess whether these interactions were a non-specific result of hyperosmotic stress-mediated molecular crowding, we examined the proximal interactions of the P-body marker DCP1A. We did not find significant enrichment of microtubule-associated proteins in DCP1A hyperosmotic stress BioID data, suggesting this change is unique to stress granule baits (Figure S3C, Table S3).

### Quantitative Analysis of Stress BioID

#### A. Identification of High-Confidence Interactions using SAINTq

Under the initial DDA-based analytical framework, a large portion of the dataset was marked by missing values across different stress time points, likely due to the semi-stochastic nature of DDA mass spectrometry (Figure S4A). To improve the quantitative analysis, we re-analyzed our BioID samples using <u>D</u>ata-Independent <u>A</u>cquisition (DIA) mode in the mass spectrometer. We built a quantitative DIA analysis workflow tailored to BioID samples to accommodate the frequent variability in interactor abundance and identity among different baits. In brief, DIA output was deconvoluted with DIA-NN^81^ for protein identification and quantification, analyzed with SAINTq^82^ to define high-confidence interactors, and linear-mixed model ANOVA to identify significant quantitative changes (Figure 2A; Figure S4B; see Methods).

**Figure 2.**
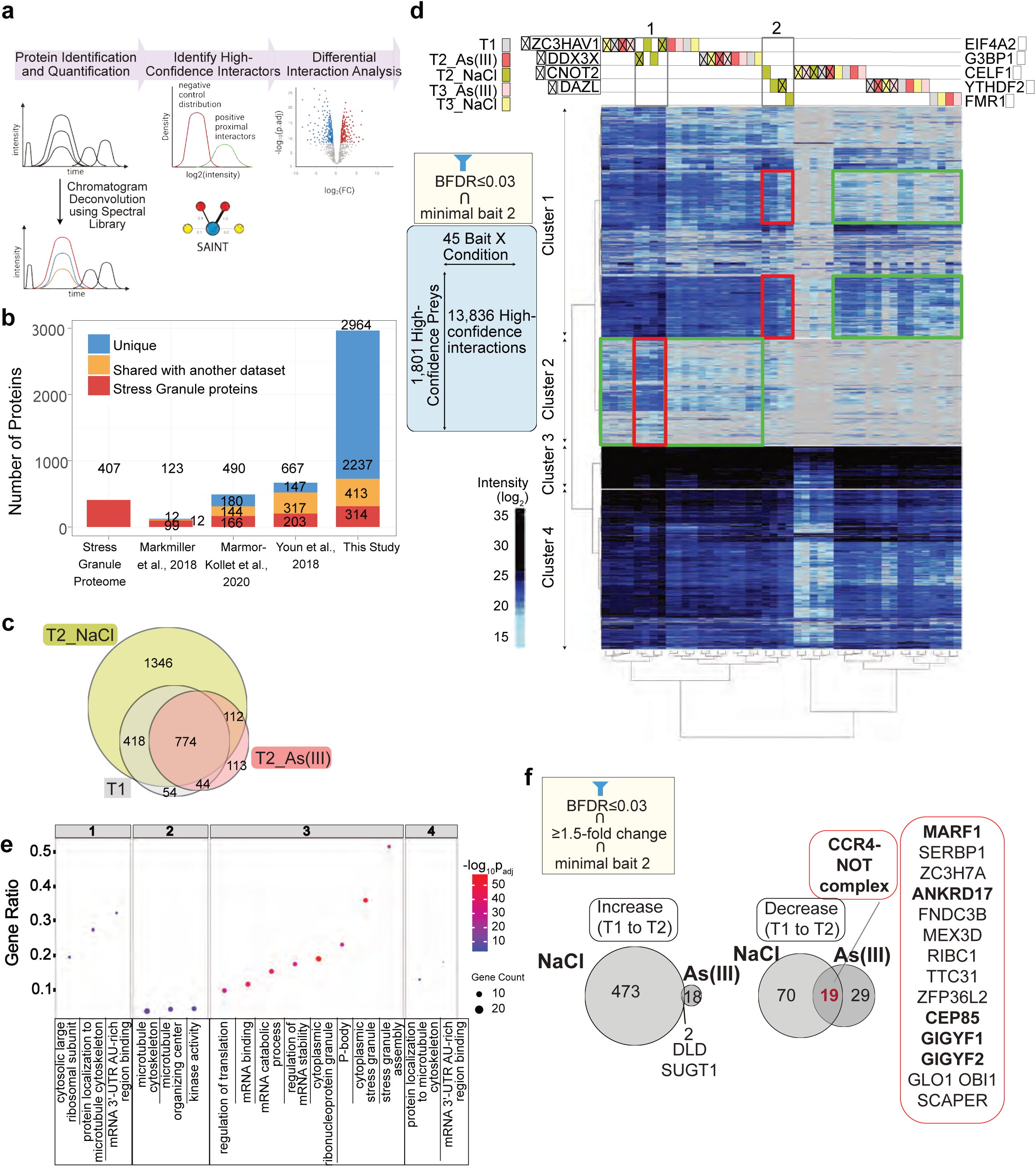
Dynamic proteomic landscapes of stress granules. (**a**) Diagram depicting stepwise DIA data analysis pipeline to improve identification and quantification. (**b**) Bar graph shows comparison to other existing datasets. Colours indicate overlap with literature-curated stress granule proteins (red), shared with another dataset but not annotated as stress granule proteins (orange), and unique to each dataset (blue). Youn *et al.,* 2018 dataset includes high-confidence proximal interactors identified by 58 stress granule proteins under no stress condition. (**c**) Venn diagram summarizes the overlap between high-confidence interactors identified in no stress (T1), T2 in oxidative stress (T2_As(III)), and hyperosmotic stress (T2_NaCl) by one or more baits. (**d**) Heatmap display of hierarchical clustering analysis of statistically significant proximal interactors identified by at least two baits in stress BioID DIA data (BFDR≤0.03). Baits and conditions are clustered on the x-axis, and high-confident interactors are clustered on the y-axis based on their abundance. Boxed areas in red highlight proximal interactions that drive clustering of baits in T2_NaCl condition compared to other conditions in green. (**e**) Dot Plot visualizes the Gene Ontology terms enriched in each cluster indicated in panel (d). (**f**) Venn diagram of high-confidence interactors displaying ≥1.5-fold change in two baits or more from no stress (T1) to stress condition (T2).

With DIA, we improved detection sensitivity and data completeness (Figure S4C-D). Biological replicates showed high reproducibility, with correlation coefficients ranging from 0.853 to 0.963 (Figure S5). After SAINTq analysis, we identified over 2,000 novel interactors compared to previously reported stress granule datasets generated using proximity-dependent biotinylation techniques (Figure 2B). Separating high-confidence interactors based on the stress condition revealed that the majority of the novel interactors are identified in the hyperosmotic stress condition (T2_NaCl). Comparisons of high-confidence interactors (BFDR ≤ 0.03, SAINTq) identified in no stress (T1) against those in oxidative (T2_As(III)) or hyperosmotic stress (T2_NaCl) show a two-fold increase in interactors unique to T2_NaCl (≥1 bait; Figure 2C). These results indicate that the dynamic rearrangements in the stress granule proximal interaction networks depend on the stress context.

To accurately discern stress granule-relevant interactions, we filtered for high-confidence interactors identified by multiple stress granule baits (BFDR ≤ 0.03; ≥ 2 baits). Under these criteria, DIA-SAINTq analysis identified 13,836 high-confidence bait-prey interactions across the entire BioID dataset, comprising 1,801 unique preys (Table S4). This yields a bait-specific recovery rate of 10-64% of the protein-protein interactions annotated in BioGRID, with stress time points observing marginally greater recovery (Table S4).

The BioID data is well visualized by hierarchical clustering analysis of the bait-condition and prey abundance matrix (Figure 2D; see Methods). Bait-condition combinations primarily clustered according to bait protein, rather than stress condition, indicating that each bait maintains a distinct labeling profile regardless of state changes. However, all baits (except CNOT2) resolve into two distinct clusters during hyperosmotic stress (top gray boxes 1 and 2 in Figure 2D). This clustering of T2_NaCl samples is attributed to proximal interactors exhibiting greater abundances (red boxes) compared to their T1, T2_As(III), or T3 sample analogs (green boxes; Figure 2D).

High-confidence preys cluster into four large groups characterized by distinct GO term enrichment (Figure 2E, Table S5). Cluster 3 preys are most abundant and comprise known stress granule components. Expectedly, these are enriched for terms including “cytoplasmic ribonucleoprotein granule” (GO:0036464) and “stress granule assembly” (GO:0034063). Cluster 1 is similarly defined by GO terms relevant to stress granule biology, such as “mRNA 3’ UTR binding” (GO:0003730), but also unexpected terms like “cytoplasmic large ribosomal subunit” (GO:0022625). An enrichment for proteins associated with the “microtubule cytoskeleton” (GO:0015630) and those presenting “kinase activity” (GO:0016301) are found in cluster 2. Preys in this cluster are detected at a higher abundance in T2_NaCl samples. Finally, cluster 4 shows enrichment for proteins in the “cytoskeleton” (GO:0005856) and “mRNA 3’-UTR AU-rich region binding” (GO:0035925). Many unanticipated GO terms belong to hyperosmotic stress-specific interactors, highlighting the context-specific nature of the stress granule proximal interaction network.

#### B. Quantitative Analysis Shows Context-Specific Rewiring of Proximal Interaction Networks During Stress Granule Assembly and Dissolution

Using the intensity-based quantitative values measured by DIA analysis, we sought to identify significant changes in the abundance of high-confidence interactors for each bait throughout the stress time course. Quantitative changes in proximal interactions for each bait-prey (e.g., FMR1-BICC1) between all time points (T1 *vs.* T2, T2 *vs.* T3, T1 *vs.* T3) were assessed using a linear mixed model ANOVA with post-hoc pairwise comparisons (Figures 2A, S4B, Methods). We analyzed 22,899 high-confidence bait-prey interactions during oxidative and hyperosmotic stress transitions, respectively (preys identified with BFDR ≤ 0.03 in any given time point for each stress condition by one or more baits). Significant changes were identified in 869 (3.8%) and 4,881 (21%) bait-prey pairs during oxidative and hyperosmotic stress and recovery transitions, respectively (≥ 1.5-fold intensity change; post-hoc p-value_adj_ ≤ 0.01; Table S6). To ensure that the observed changes in the BioID data were not due to global proteome changes during stress, we induced oxidative and hyperosmotic stresses in the G3BP1 Flp-in HEK293 cell line and analyzed the total proteome using the quantitative DIA analysis workflow (see Methods). Differential analysis of global protein abundance revealed minimal changes between no stress and each stress condition, indicating that observed abundance changes are due to proximal interaction changes, not due to proteome abundance changes upon stress (Figure S6A; Table S7).

Despite the apparent increase in protein concentration within stress granules, contrary to our expectations, high-confidence prey abundance remained largely unchanged following exposure to oxidative stress (T1 to T2; Figure S6B). In contrast, the corresponding distribution tended toward positive fold-change during hyperosmotic stress. Interactions that increased during hyperosmotic stress would later return to no stress-state levels, as indicated by the negative fold-changes observed during the transition from stress to recovery (T2 to T3). These increased interactions during hyperosmotic stress were specific to high-confidence interactors, with the overall intensity of non-significant interactions remaining similar between stress and no stress state (median fold-change for non-significant interactors: 1.05 *vs.* high-confidence interactors: 1.27). This suggests increases in stress granule-specific interactions rather than as a result of general overcrowding in the cytoplasmic space.

#### C. Validation of Identified Stress-Specific Interactors

We selected several proteins that increased in a stress-specific manner during stress granule assembly for colocalization analysis with the stress granule marker G3BP1 for validation. TRIM56, BICC1, LARP6, and NYNRIN increased in abundance during oxidative stress, but remained unchanged during hyperosmotic stress. Colocalization analysis was performed under both stress conditions in HeLa cells using antibodies against endogenous proteins or by stable expression of mCherry-tagged proteins. All tested preys localized to stress granules regardless of the inducing agent, except for TRIM56. TRIM56 recruitment to stress granules was specific to oxidative stress (Figure S6C; data not shown). BICC1, LARP6, and NYNRIN localized to stress granules during hyperosmotic stress, despite their unchanged abundance measurements. This is not unexpected, as the majority of known stress granule proteins displayed minimal changes in abundance under either stress condition. We further investigated whether TRIM56 plays a role in stress granule assembly. The loss of TRIM56 through CRISPR-mediated knockout did not show defects in stress granule assembly, indicating that increased association with specific components does not entail their essentiality for stress granule assembly (data not shown).

Next, we tested if hyperosmotic stress-specific interactors—60S large ribosomal subunits and centrosome-associated proteins—localize to stress granules induced by hyperosmotic stress. When we tested the localization of RPL32 and RPL36 using immunofluorescence microscopy, their cytoplasm-wide diffusive signal made it difficult to assess their colocalization. To reduce their diffusive signal in the cytoplasm, we permeabilized the plasma membrane with a low concentration of digitonin before fixation^83^ (see Methods). The digitonin-permeabilized immunofluorescence images showed partial colocalization of G3BP1 with RPL32 or RPL36 subunits in stress granules (Figure S7A-B), consistent with a recent study reporting the presence of 80S ribosomes in stress granules using cryogenic correlative light and electron microscopy^84^. The extent of colocalization between cytoplasmic RPL32 *vs.* G3BP1 or cytoplasmic RPL36 *vs.* G3BP1 was consistently the greatest in hyperosmotic stress compared to untreated or oxidative stress conditions (Figure S7C-D), corroborating our stress BioID results. While the trend showed good agreement, the extent of colocalization in hyperosmotic stress was significantly higher than the untreated condition only for RPL36, not for RPL32. As for colocalization analysis of a subset of centrosome-associated proteins (CEP170B, PCNT and CEP192), we found poor enrichment within stress granules during hyperosmotic stress (data not shown). We attribute this to their proximal interactions occurring in the dilute phase.

In summary, we validate TRIM56 as an oxidative stress-specific and RPL36, a large ribosomal subunit, as a hyperosmotic stress-specific interactor for stress granules. Further testing is required to elucidate the nature of stress granule interactions with microtubule- and centrosome-associated proteins.

### Dynamic Proximal Interaction with the CCR4-NOT Complex is Universal during Stress Granule Assembly

We hypothesized that quantitative changes across multiple baits and common to both stressors could help identify a shared mechanism underlying stress granule assembly or stress granules’ shared function in cellular adaptation to stress. To assess condensate-level changes for each stress, proximal interactions were filtered for those demonstrating consistent trends in their interaction changes (≥ 1.5-fold increase, decrease or unchanged) across two or more baits (Table S6). Consistent changes were observed during hyperosmotic stress (n=564) and oxidative stress (n=68).

Next, we assessed shared changes in abundance in both stressors (Figure 2F). While only two high-confidence preys (SUGT, DLD) increased in both oxidative and hyperosmotic stress (≥1.5-fold; ≥2 baits), a significant amount of overlap was observed among interactors that consistently decreased in abundance (30% and 48% of decreasing preys in hyperosmotic and oxidative stress, respectively). These included RNA-binding proteins involved in mRNA deadenylation, translation repression and mRNA decay (*e.g.,* CCR4-NOT complex components and interactors, including TOB1, EIF4E2, and EIF4ENIF1).

GO term enrichment analysis of interactors that shift in abundance during stress and recovery (Table S6) identified terms related to ‘CCR4-NOT deadenylase complex’ among preys that decrease during stress granule assembly and later increase as they disassemble. This trend is consistent across both conditions, as evidenced by the quantitative changes in G3BP1 proximal interactors (Figure S8A). Notably, the decreased association with CCR4-NOT components from T1 to T2 quickly recovers during the T2 to T3 transition (Table S6). No changes in total protein abundance of the CCR4-NOT components were observed (Table S7). Together, these observations show the coupling of CCR4-NOT proximal interactions with stress granule assembly dynamics, with no changes in CCR4-NOT abundance.

### Loss of the CCR4-NOT complex Blocks Lipoamide-induced Inhibition of Stress Granule Assembly

As stated, pre-treatment with lipoamide effectively inhibits stress granule assembly (Figure 1C). To further characterize this inhibitory pathway and elucidate mechanisms regulating stress granule assembly, we performed a genome-wide CRISPR-Cas9-mediated gene knockout screen (Figure 3A). HeLa cells stably expressing Cas9 and single guide RNAs (sgRNAs) were pre-treated with lipoamide prior to stress with potassium arsenate (As (V)). Confocal images of stress granules, identified by punctate forms of stably expressed FUS-GFP signals, were segmented, quantified, normalized, and transformed into z-scores (Figure 3B; Methods). Notably, the knockout of SRSF1 abolished the inhibitory effects of lipoamide (z-score = 17.5; Figure 3B). This result substantiates the screen, as SRSF1 has previously been shown to prevent lipoamide activity on stress granules^67^. Other gene knockouts (*e.g.,* SLC3A2, ABCC1, large ribosomal subunit proteins, and PPIP5K2) also produced similar inhibitory effects.

**Figure 3.**
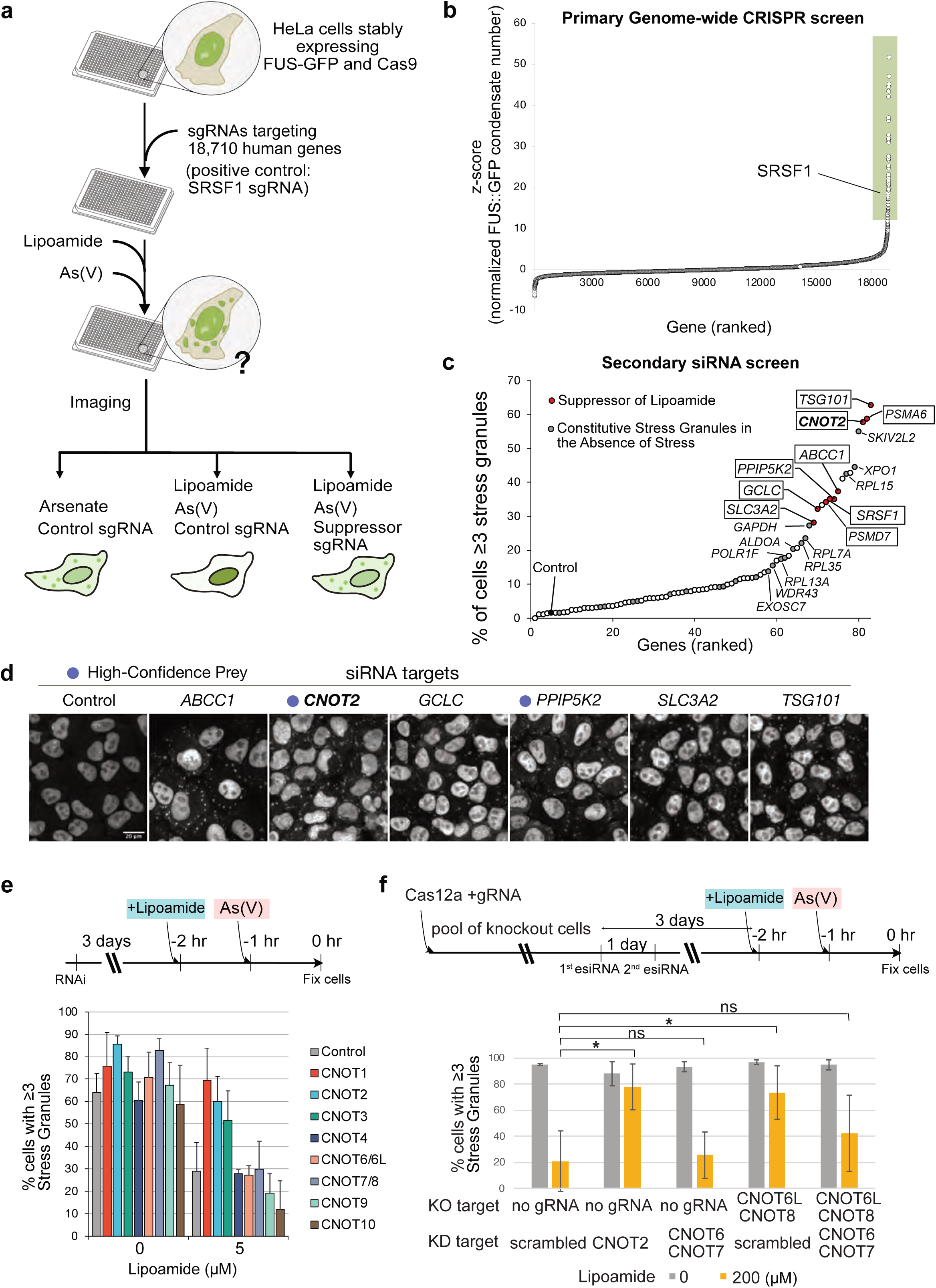
A genome-wide lipoamide suppressor screen identifies components of the CCR4-NOT complex to bypass the lipoamide effect on stress granules. (**a**) Schematics showing genome-wide CRISPR Cas knock-out screen to identify genes required for lipoamide inhibitory activity on stress granule formation. (**b**) A rank-ordered visualization of candidate genes whose knock-out results in suppression of lipoamide activity, including its known target, SRSF1. (**c**) Secondary siRNA screen and their results. Lipoamide suppressors validated in oxidative stress condition are indicated in a box (filled in red). Targeting a subset of genes resulted in constitutive induction of stress granules in the absence of stress (filled in gray; no box). (**d**) Representative images of validated lipoamide suppressors. Magenta dot: high-confidence interactors in this stress BioID dataset. Scale bar = 20 µM. (**e**) Examination of individual CCR4-NOT complex components on their requirements for lipoamide activity on stress granules using siRNA-mediated knockdown in HeLa cells (top). Cells were pretreated with 5 µM of lipoamide, then stressed with 1 mM of potassium arsenate (As(V)) (additional lipoamide concentration data available in Supporting Data). Quantification of the percentage of cells with three or more cytosolic FUS-GFP puncta is plotted against varying lipoamide pre-treatment conditions from three biological replicates of siRNA experiments (bottom). (**f**) The effect of combined loss of enzymatic subunits via knockout or knockdown on lipoamide activity on stress granules in HeLa FUS-GFP cells (top). Cells were pretreated with 200 µM of lipoamide, then stressed with 20 mM of sodium arsenate (As(V)). The percentage of cells with three or more cytosolic FUS-GFP puncta is plotted using quantifications from two biological replicates of experiments (bottom; * p-value<0.05 t-test).

The microscopy data were manually reviewed, and 81 genes were selected for further validation using endoribonuclease-prepared small interfering RNA (esiRNA)-mediated gene knockdown. Using the same experimental workflow and analysis pipeline as the primary screen, we observed candidates that robustly induced stress granule assembly in cells pre-treated with lipoamide (Figure 3C). Examination of stress granule dynamics in the absence of lipoamide, arsenate (As(V)), or both revealed that the depletion of 22 genes induced constitutive stress granule assembly (filled in gray, Figure 3C). These genes, including those that encode helicases, exosome components, ribosome components, and components involved in ribosome biogenesis (RNA polymerase I and WDR43), induced cytosolic FUS-GFP puncta in the absence of cellular stress. Excluding these genes, the depletion of nine others effectively blocked the inhibitory effect of lipoamide on stress granule assembly, including our positive control, SRSF1 (genes in boxes; Figure 3C).

Among these, GCLC and SLC3A2 are involved in glutathione biosynthesis^85^, supporting the redox-associated role of lipoamide for stress granule dynamics^67^. Additionally, the proteasome subunits like PSMA6 and PSMD7 were validated, consistent with prior observations of proteasomal involvement in stress granule dynamics^86^. Notably, PPIP5K2, a bifunctional inositol kinase, and CNOT2, a component of the CCR4-NOT complex, were both identified as high-confidence interactors in the stress BioID data (Table S4).

Given the dynamic changes in proximal associations between the CCR4-NOT complex components and stress granule baits, we focused our further investigation on CNOT2. Other CCR4-NOT complex components were depleted to determine whether CNOT2 itself or the complex plays a role in stress granule dynamics. We depleted CCR4-NOT complex components individually or in pairs of the deadenylase paralogs using siRNA (*i.e.,* CNOT7 and CNOT8 or CNOT6 and CNOT6L). Depletion of the NOT module proteins abrogated lipoamide-mediated inhibition of stress granule assembly, whereas depleting the paralog pairs CNOT7-CNOT8 or CNOT6-CNOT6L did not (Figure 3E). Next, we tested removing or depleting deadenylase subunits in different combinations by employing both CRISPR Cas12a knockout and siRNA-mediated knockdown. Removing the CNOT6-CNOT7 pair yielded no suppression of the lipoamide effect, whereas loss of the CNOT6L-CNOT8 pair efficiently blocked lipoamide’s effect on stress granules (Figure 3F). Surprisingly, removing all enzymes (CNOT6, CNOT6L, CNOT7, and CNOT8) did not block lipoamide’s effect on stress granules, which may be due to the compensatory increase in protein abundance we observed for the remaining subunits. Particularly, the NOT module (CNOT1, 2 and 3) significantly increased in abundance by 1.5∼3 fold (Figure S8B). Together, this suggests two possible mechanisms at play in blocking lipoamide’s effect: one that depends on the deadenylase activity and another that relies on deadenylase-independent function(s) of the NOT module.

### Regulation of Poly(A) Tail Lengths is Important for Stress Granule Assembly

We considered two scenarios in which loss of the CCR4-NOT complex could block lipoamide’s effects. First, lipoamide may regulate CCR4-NOT complex function to influence stress granule assembly. However, previous thermal proteome profiling assays using lipoamide showed no change in melting temperature for CCR4-NOT complex components (CNOT1, 2, 3, 4, 6, 7, 10 and 11), advocating against their direct interaction^67^.

Alternatively, CCR4-NOT may operate in a parallel pathway, distinct from lipoamide regulation, to promote stress granule assembly. Given the established relationship between poly(A) tail length and stress granule assembly ^61^, and the broader role of mRNAs in condensate assembly ^87-91^, we hypothesized that poly(A) lengths regulated by the CCR4-NOT complex could modulate stress granule assembly. Longer poly(A) tails may enhance scaffolding for stress granule assembly by providing additional binding sites for poly(A) binding protein (*i.e.,* PABPC1), whereas shorter poly(A) tails could diminish this ability.

To assess the influence of poly(A) tail shortening on stress granule assembly, we utilized cordycepin, an adenosine analog that inhibits polyadenylation, to reduce global poly(A) tail length^92^. Cordycepin pre-treatment prior to oxidative stress significantly reduced stress granule and P-body numbers, marked by G3BP1 and DCP1A, in a concentration-dependent manner (Figures 4A-B). PABPC1 recruitment to stress granules was likewise reduced in a dose-dependent fashion, consistent with shorter poly(A) tail diminishing PABPC1-mRNA binding (Figures S8C-D). This can be contrasted with our earlier genetic screens, where inhibiting deadenylase activity by reducing the NOT module of the CCR4-NOT complex positively influenced stress granule assembly. Specifically, in the absence of lipoamide, the knockdown of CNOT1, CNOT2, and CNOT7/8 significantly increased the number of stress granules following oxidative stress (Figure 4C). Together with prior studies reporting global poly(A) tail lengthening following the loss of catalytic enzymes (CNOT7/8) and the NOT module (CNOT1 and CNOT2)^45,93,94^, we propose that poly(A) lengthening promotes stress granule assembly.

**Figure 4.**
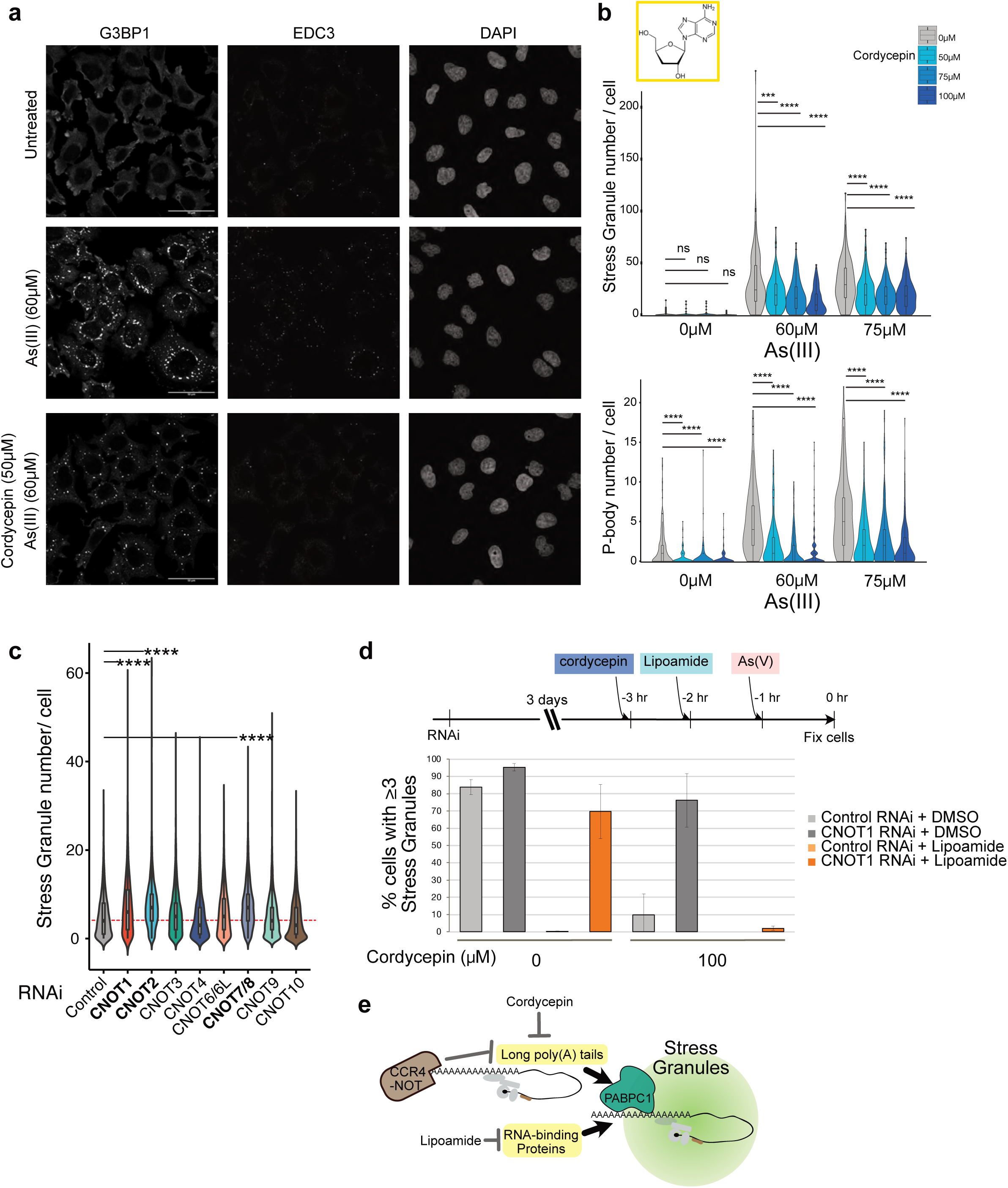
Altering Poly(A) lengths affects efficient stress granule assembly. (**a**) Representative images show the effect of cordycepin (50 µM, 2 h) on stress granules (G3BP1) and P-bodies (EDC3) during oxidative stress (sodium arsenite, As(III); 60 µM, 1 h) in HeLaFlp-In T-REx cells. Scale bar = 50 µM. (**b**) Violin-Box plot of stress granule and P-body numbers in HeLa Flp-In T-REx cells pretreated with increasing concentrations of cordycepin for 2 h, followed by no treatment or combined with sodium arsenite, As(III) stress (60 µM or 75 µM, 1 h). Quantifications from three biological replicates were assessed with Mann Whitney test for statistical significance compared to 0 cordycepin treatment (*** p_adj_ < 10^-3^, **** p_adj_ < 10^-4^). (**c**) Violin-Box plot shows the effect of loss of specific CCR4-NOT components on stress granule numbers (cytoplasmic FUS-GFP) during oxidative stress (1mM sodium arsenate, As(V), 1 h). The number of stress granules in siRNA-treated samples *vs.* control tested from three biological replicates of siRNA experiments is assessed for statistical significance using the Mann-Whitney test (**** p_adj_ < 10^-4^). Red dotted line: the median stress granule number in control. (**d**) Bar graph visualization of the effect of CNOT1 knockdown on stress granule formation during oxidative stress (1mM potassium arsenate, As(V)) in HeLa cells pretreated with conditions that inhibit stress granule formation (cordycepin, 10 µM lipoamide, or cordycepin and 10 µM lipoamide combined). The sequence of treatments is shown in the top panel. The bar graph depicts data from three biological replicates, quantified from over 300 cells. (**e**) Model of two paralleled mechanisms driving stress granule formation. Lengthening global poly(A) tails promotes stress granule formation by providing longer RNAs for PABPC1 scaffolding. The CCR4-NOT complex and cordycepin shorten Poly(A) tails, preventing efficient stress granule formation. Lipoamide blocks the oxidation of RNA-binding proteins that promote stress granule formation^56^.

We then combined siRNA-mediated knockdown of the CCR4-NOT complex with cordycepin, lipoamide or both to assess whether poly(A) lengthening can counteract the stress granule-inhibitory effects of these treatments during oxidative stress. While cordycepin pre-treatment alone significantly reduced stress granule assembly, knockdown of CNOT1 prior to acute cordycepin treatment induced stress granule assembly (Figure 4D). This suggests that the stimulatory effects of CCR4-NOT knockdown (*i.e.,* global poly(A) lengthening) can negate the inhibitory effects of acute cordycepin treatment. However, CNOT1 knockdown did not overcome the combined inhibitory effects of both lipoamide and cordycepin, leading to defective stress granule assembly. This suggests two parallel pathways regulating stress granule assembly: one through redox regulation of stress granule proteins (via lipoamide) and the other through control of poly(A) tail lengths (via cordycepin and CCR4-NOT activity) (Figure 4E).

### CCR4-NOT complex is Sequestered to Stress Granules During Stress

Components of the CCR4-NOT have been shown to co-localize with P-body markers in overexpression systems^95,96^. Under endogenous conditions, we observed poor colocalization of CNOT1 with the P-body markers EDC3 (at their endogenous levels) or with DCP1A (stably and inducibly expressed under no stress condition) in HeLa and HEK293 Flp-In T-REx cells (Figures 5A, S9A). However, under both oxidative and hyperosmotic stress, CNOT1 formed large condensates adjacent to P-bodies, similar to the docking behaviour of stress granules^97^. When directly tested, CNOT1 colocalized with stress granule marker G3BP1 during both oxidative and hyperosmotic stress in HeLa and HEK293 cells (Figures 5B, S9B). Other CCR4-NOT components, including the NOT module (CNOT2) and the deadenylase subunits, CNOT7 and CNOT8, localized to stress granules under these conditions in HeLa and HEK293 cells (Figures 5B, S9B). Stress-induced localization of CCR4-NOT complex components to stress granules was further confirmed by inducible expression of tagged proteins: 3x-FLAG-CNOT2 in HEK293 and mNeon-tagged CNOT2, CNOT6L, or CNOT7 in HeLa cells expressing mScarlet-G3BP1 (Figures S9C-D). To determine whether the CCR4-NOT complex remains intact during partitioning to stress granules, we visualized CNOT2 BioID interactions with other complex subunits. We observed sustained proximal interactions between CNOT2 and the rest of the subunits upon stress, indicating that the complex remains intact in stress granules (Figure 5C). Furthermore, high-resolution imaging of CNOT1 and G3BP1 during oxidative stress revealed distinct patterns of CNOT1 enrichment within the G3BP1-positive stress granules, indicating an organized partitioning of the complex into nano-compartments within stress granules (Figure 5D).

**Figure 5.**
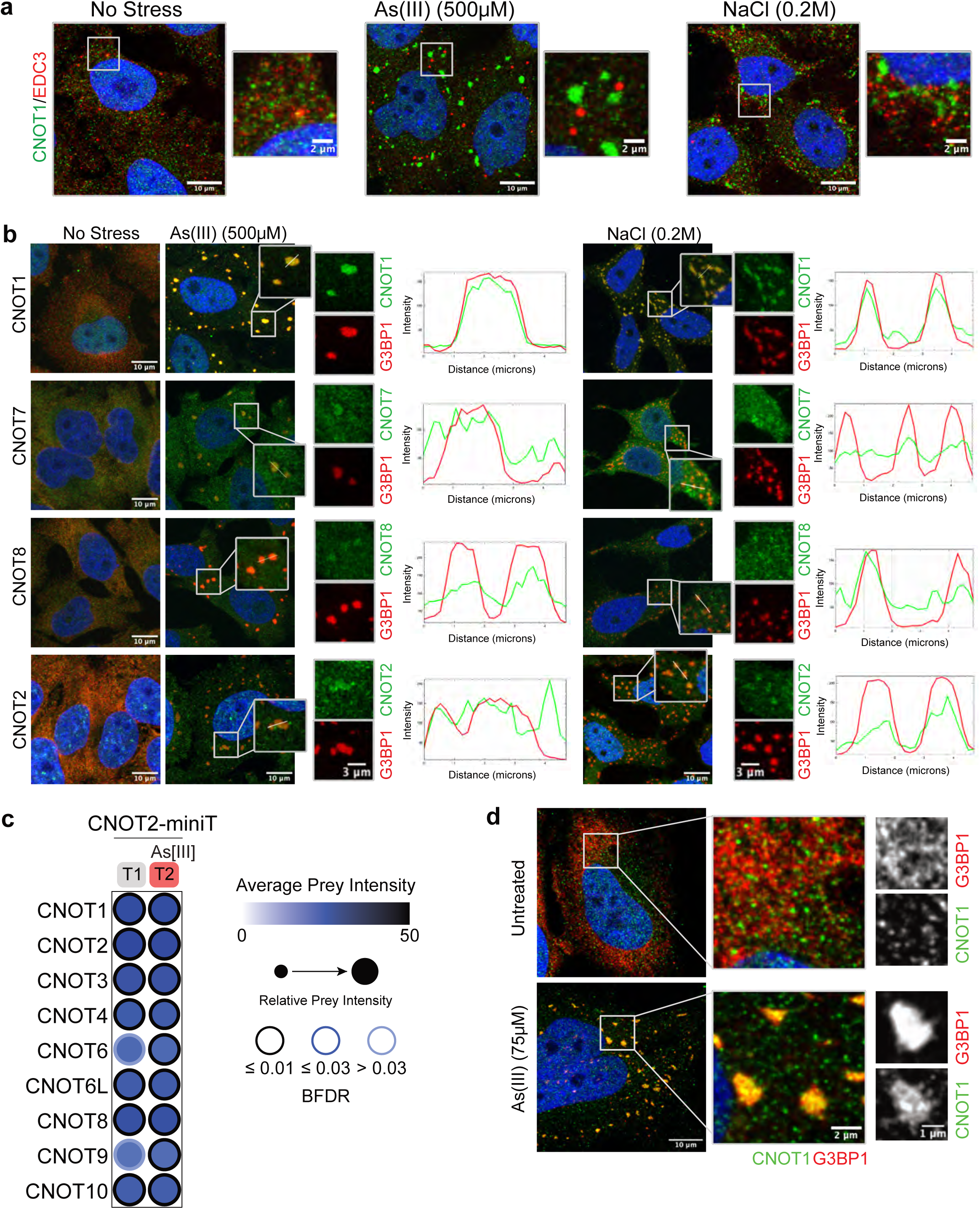
Stress Granule localization of the CCR4-NOT complex. (**a**) Immunofluorescence microscopy of CNOT1 (green), P-body marker EDC3 (red), and nuclei (DAPI) in HeLa cells at steady state and stress conditions (500 µM sodium arsenite, As(III), 30 min or 0.2 M NaCl, 1 h). Scale bar = 10 µM. (**b**) Immunofluorescence microscopy of CNOT1, CNOT2, CNOT7 and CNOT8 (green), stress granule marker G3BP1 (red), and nuclei (DAPI) during oxidative (sodium arsenite, As(III), 500 µM, 30 min) and hyperosmotic stress (sodium chloride, 0.2 M, 1 h) in HeLa cells. Line plots represent the fluorescent intensity included in the selected insets. Scale bars = 10 µM and 3 µM (from left to right zoomed insets). (**c**) Dot plot visualization of CNOT2 proximal interactions with the rest of the CCR4-NOT complex subunits during no stress (T1) and oxidative stress (T2_As(III)). (**d**) Lightning mode confocal microscopy of CNOT1 (green), stress granule marker G3BP1 (red), and nuclei (DAPI) during oxidative stress (sodium arsenite, As(III), 75 µM, 1 h) and at steady state in HeLa cells. Scale bars = 10, 2 and 1 µM (from left to right zoomed insets).

CNOT1 has been reported as a component of the assemblysomes, formed by translationally inactive ribosome-nascent chain complexes during proteotoxic stress^98^. To distinguish if stress-induced CNOT1 puncta are assemblysomes or not, we tested a characteristic that distinguishes the assemblysomes from stress granules and P-bodies: resistance to cycloheximide^99^. Arsenite (As(III)) stress-induced CNOT1 puncta colocalizing with G3BP1 dissolved upon cycloheximide treatment, indicating that CNOT1 puncta are not assemblysomes (Figure S10). Together, these findings demonstrate that, endogenous CCR4-NOT complex shows poor P-body localization and the complex is robustly recruited to stress granules during cellular stress.

### CCR4-NOT Complex Sequestration in Stress Granules Promotes Poly(A) Lengthening of Stress-Survival Transcripts

The CCR4-NOT complex promotes mRNA decay and translation repression via shortening poly(A) tails of transcripts^44,45,100^. To test if stress granule-localization of CCR4-NOT complex was responsible for poly(A) tail lengthening^56-59^ observed in stress, we blocked stress granule assembly using lipoamide, and investigated the effect on poly(A) tails and abundance of the transcriptome. We collected total RNAs from HeLa cells under three conditions: no stress, sodium arsenate (20mM As(V)) stress, and lipoamide pre-treatment followed by sodium arsenate stress (200 μM lipoamide + 20mM As(V)) during which lipoamide was sustained (Figure 6A). Lipoamide blocked stress granule assembly and released the CCR4-NOT scaffold protein, CNOT1, back to the cellular environment during cellular stress (Figure S11A). We utilized Nanopore Direct RNA Sequencing (DRS) to determine the poly(A) tail lengths and the abundance of transcriptome in each condition (n=3; see Methods).

**Figure 6.**
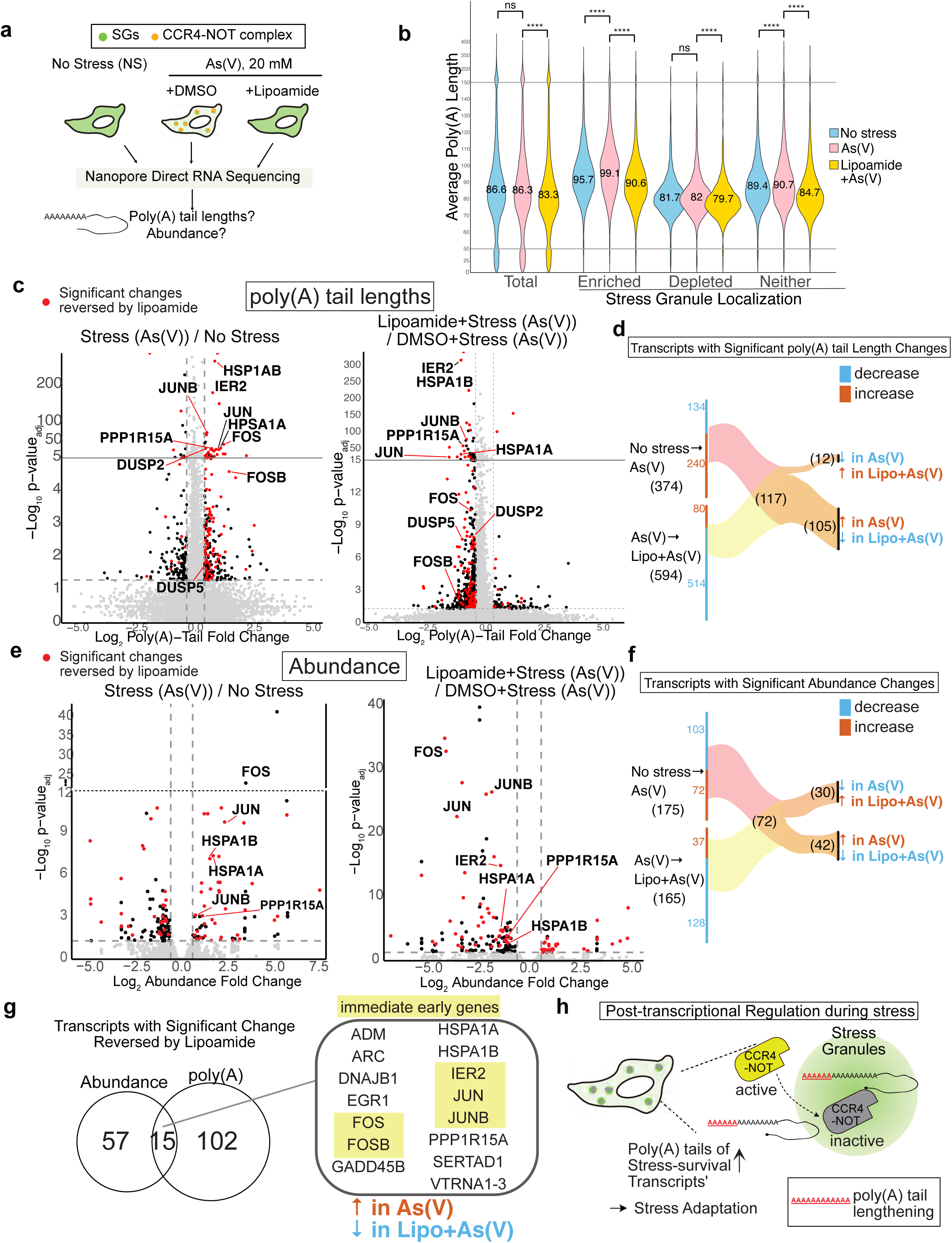
Nanopore direct RNA sequencing demonstrates that sequestration of CCR4–NOT into stress granules promotes poly(A) tail lengthening of stress-survival transcripts. (**a**) Schematic showing experimental design for the transcriptomics analysis performed using Nanopore direct RNA sequencing (DRS). Abundance and poly(A) length measurements were extracted from the direct RNA sequencing data. (**b**) Transcriptome-wide distribution of gene-level average poly(A) lengths during no stress, stress and lipoamide+stress, with the average values indicated in the middle of each violin plot. Groups indicated the entire transcriptome (left), or subgroups of transcripts categorized based on their enrichment in stress granules (right). Mann-Whitney U test on three biological replicates was performed with Bonferroni correction to determine significant differences in poly(A) length distributions (** padj <10-2, **** padj <10-4). (**c**) Volcano plot of poly(A) length fold-changes from no stress to stress (As(V)), and stress (As(V)) to lipoamide + stress (As(V)) and associated adjusted p-values as determined by TAILCaller and subsequent post-hoc tests. A >|1.3|-fold cut-off for fold-change was used (n=3). (**d**) Sankey plot shows the distribution of transcripts whose poly(A) tail lengths were significantly changed in both condition-condition comparisons and their direction of change. (**e**) Volcano plot of abundance fold-changes from no stress to stress (As(V)), and stress (As(V)) to lipoamide + stress (As(V)), and associated adjusted p-values, as determined by DESeq2. A >|1.5|-fold cut-off for fold-change was used (n=3). (**f**) Sankey plot shows the distribution of transcripts whose abundances were significantly changed in both condition-condition comparisons and their direction of change. (**g**) A Venn diagram shows genes whose significant changes in either poly(A) lengths or abundance upon stress were reversed in lipoamide+stress conditions. The 15 shared genes all show stress-induced increases in poly(A) tail length and transcript abundance, changes that are fully reversed by lipoamide treatment. (**h**) Schematics show a proposed model for stress granule function in post-transcriptional regulation. Upon stress, the CCR4-NOT complex is sequestered to stress granules, resulting in poly(A) tail lengthening of stress-survival transcripts, promoting stress adaptation.

To examine the effect of stress and blocking stress granules on poly(A) tail lengths, we calculated the average poly(A) tail lengths at the gene level and visualized their distribution. Globally, the average poly(A) lengths minimally changed between no stress and stress conditions. However, lipoamide-mediated inhibition of stress granule assembly reduced the average poly(A) tail lengths compared to the stress condition, which we attribute to the active CCR4-NOT complex that failed to be sequestered to stress granules (Figure 6B). Reduced poly(A) effect is not due to lipoamide agent on its own, as lipoamide treatment in the absence of stress caused a moderate increase in poly(A) tails (Figure S11B).

Furthermore, grouping transcripts based on their stress granule localization^30^ revealed biases in stress-induced poly(A) tail lengthening. Transcripts enriched in stress granules (‘enriched’) or neither enriched nor depleted (‘neither’) showed statistically significant increases in poly(A) length upon stress. These differences suggest that transcripts enriched in stress granules, or neither enriched nor depleted, are more affected by CCR4-NOT’s stress granule localization. This corroborates the idea that CCR4-NOT deadenylase activity depends on the translation efficiency of the transcripts^44^; the complex rapidly deadenylates transcripts with poor translation efficiency, which is a characteristic of stress granule-enriched transcriptome^30^. Regardless of transcript localization, average poly(A) tails were shortened when lipoamide blocked CCR4-NOT sequestration into stress granules compared to the stress condition (Figure 6B).

To identify genes significantly regulated by CCR4-NOT complex sequestration to stress granules, we employed the TAILcaller^101^, which determines significant poly(A) tail distribution changes in two or more conditions (Figure S11C and Methods). Analyzing reproducible changes in poly(A) lengths between no stress vs. stress (As(V)) or stress (As(V)) vs. lipoamide+stress (As(V)) revealed 374 transcripts and 594 transcripts with significant poly(A) length distribution changes, respectively (p_adj_ <0.05 and >|1.3|-fold in median lengths, Figure 6C, Table S8). Consistent with the global trend, the majority of transcripts with significant poly(A) tail changes from the no stress to stress comparison showed an increase, while poly(A) tails generally decreased between stress *vs.* lipoamide+stress comparison, reversing the effect of stress on poly(A) tail lengths (Figure 6D). In other words, poly(A) tail lengths in the lipoamide + stress condition were similar to the no stress condition (Figure S11D, left panel).

Analysis of significant abundance changes under the same conditions using DESeq2^102^ showed the same reversible trend. Pairwise comparisons of no stress vs. stress and stress vs. lipoamide+stress revealed 175 and 165 transcripts, respectively, with significant abundance changes, showing reversal of stress-induced changes in lipoamide+stress condition (Figure 6E). Focusing on stress-induced changes reversed by lipoamide-mediated inhibition, we found a total of 72 transcripts showed significant changes in either direction; 42 significantly increased in abundance upon stress (AsV), which decreased back to the stress level in the lipoamide+stress condition, and 30 showed the opposite trend (Figure 6F).

Comparison of transcripts with significant poly(A) tail and abundance changes that are reversed by blocking stress granule assembly identified 15 that are shared, all of whose poly(A) tails and abundance increased upon stress and decreased in the lipoamide+stress condition (Figure 6G). The shared group includes PPP1R15A (GADD34), the regulatory component of the protein phosphatase PP1 complex responsible for inactivating the integrated stress response by dephosphorylating p-eIF2α^103^, chaperones (HSPA1A and HSPA1B), and immediate early genes (FOS, FOSB, IER2, JUN and JUNB) activated by the MAPK signaling pathways (ERK1/2, p38 and JNK1)^104^, all of which play roles in stress adaptation and promotion of survival (Figure 6G). These genes underlie the specific GO terms observed for this shared group (*e.g.,* genes involved in the regulation of cell cycle and cellular response to oxidative stress; Figure S12A, Table S8). Visualizing poly(A) lengths and abundances of individual transcripts in the shared group shows a clear correspondence between poly(A) tails and abundance, which was not seen in the non-significant transcripts (Figure S12B-C). Abundance changes for 4 of the shared group were further validated using RT-qPCR in addition to 6 transcripts showing an opposite pattern (decrease upon stress; Figures S12D-E). Given that these are steady-state measurements, we cannot distinguish if poly(A) lengthening and increased abundance observed in stress-induced survival transcripts reflect reduced decay (*i.e.,* reduced CCR4-NOT activity) or newly synthesized transcripts. Regardless, their stress-induced increase in abundance and extended poly(A) tails depend on the formation of stress granules.

Transcripts that showed significant poly(A) tail length changes, but no significant changes in abundance (*i.e.,* poly(A) changes only), were enriched for those regulating transcription by RNA polymerase II, intrinsic apoptotic signaling pathway in response to DNA damage and MAP kinase tyrosine/serine/threonine phosphatase activity (DUSP2 and DUSP5). This suggests that transcripts encoding proteins involved in transcription, apoptosis, and regulation of the MAPK signaling pathway are regulated by stress granule formation via poly(A) tail lengthening. Conversely, transcripts showing significant abundance changes, but no significant poly(A) tail length changes (*i.e.,* abundance changes only), also showed enrichment for transcription-related terms (e.g., DNA-binding transcription factor binding), while showing terms unique to this group, like regulation of cell-cell adhesion. The abundance of these transcripts is likely regulated by post-transcriptional mechanisms other than poly(A) tail lengthening.

Together, these analyses uncovered a subset of transcripts encoding proteins involved in stress response or transcription whose poly(A) tail or abundance are regulated by CCR4-NOT sequestration to stress granules.

## DISCUSSION

Biomolecular condensates participate in a wide range of biological processes, yet their dynamic nature and lack of physical boundaries (unlike membrane organelles) pose significant challenges for study. Stress granules, a condensate formed upon cellular stress, contain thousands of mRNAs and hundreds of proteins^30,31,105^. Despite their widespread association with stress response, the precise functions of stress granules remain controversial. In this study, we leveraged multi-bait BioID profiling and quantitative mass spectrometry to map the dynamic components of stress granules proteomes. Importantly, our approach integrated quantitative protein-protein interaction data with chemical genetic interaction measurements, two orthogonal sources of biological information^106,107^, to identify CCR4-NOT complex.

BioID profiling of stress granule-localized G3BP1 across a stress time course revealed noticeable changes in proximal interactions accompanying the emergence of microscopically visible stress granules (Figure 1B). Extending this approach to total of nine high-information baits allowed us to define distinct proteomic trajectories of stress granules formed under oxidative versus hyperosmotic stress. These stressors disrupt cellular homeostasis in fundamentally different ways. Oxidative stress generates excess reactive oxygen species that damage macromolecules (*e.g.,* DNA, proteins) and organelles (*e.g.,* endoplasmic reticulum and mitochondria)^73-75^, triggering the integrated stress response and stress-activated MAPK signaling pathways, leading to global translation repression, transcriptional reprogramming and metabolic remodeling. In contrast, hyperosmotic stress induces rapid water efflux within seconds, increasing intracellular macromolecular crowding and promoting condensates enriched in WNK kinases, TSC22D and NRBP proteins^77^. Condensate formation, coupled with WNK kinase activation, drives the activation of ion transporters, channels and pumps to restore cell volume through regulatory volume increase^76^.

Consistent with the distinct cellular challenges posed by these stressors, we observed context-specific proximal interaction networks of stress granules. We uncovered ∼2,000 novel proximal interactions—mostly unique to hyperosmotic stress—including many interactions with large ribosomal subunits and cytoskeletal proteins. Immunofluorescence microscopy validation revealed proteins that localize to stress granules in a context-specific manner: TRIM56 upon oxidative stress and a large ribosomal subunit, RPL36, upon hyperosmotic stress (Figures 6C, S7), consistent with our BioID results. We also detected more than 500 dynamic protein components (Figure 2F), which differed substantially between stress types, revealing striking plasticity in the stress granule proteome. Despite these differences in protein associations, stress granule transcriptomes are reported to remain highly correlated between oxidative and hyperosmotic stress^108^, suggesting that protein dynamics, rather than RNA content, may play a more prominent role in stress-specific adaptations.

A shared feature of both stress conditions was the changes in association of the CCR4-NOT complex and its interacting proteins with stress granules (Figure 2F). In parallel, an imaging-based, genome-wide CRISPR screening identified nine genes whose loss suppressed the stress granule-inhibiting effects of lipoamide. These genes include CNOT2, a key component of the CCR4-NOT deadenylases complex (Figure 3D). Loss of any NOT module protein (CNOT1, CNOT2 and CNOT3) or deadenylase subunits (CNOT6L and CNOT8) abolished lipoamide’s inhibitory effects (Figures 3E-F). Given that the CCR4-NOT complex is the principal cytoplasmic deadenylase^44-46^, we examined whether poly(A) tail length modulates stress granule assembly. Poly(A) shortening inhibited stress granule assembly and PABPC1 recruitment, whereas poly(A) lengthening had the opposite effect (Figures 4B-C, Figures S8C-D). We propose that loss of the NOT module or deadenylase subunits bypasses lipoamide activity either by increasing RNA-based intermolecular interactions^87,88,90,91,105^ or providing additional PABPC1 binding sites via poly(A) tail lengthening^61^ (Figure 4E). Together, these results reveal a functional link between mRNA deadenylation and stress granule formation.

Microscopy-based analysis demonstrated that the CCR4-NOT complex localizes to stress granules during both oxidative and hyperosmotic stress, rather than to P-bodies as previously reported (Figure 5). This was unexpected given the reduced proximal associations detected in BioID experiments (Figure 2F, Figure S8A). We attribute this discrepancy to the *cumulative* nature of BioID experiments, which report the <u>integrated sum of interactions</u> occurring inside and outside stress granules over time. We propose that overall proximity signals decrease during stress either due to reduced transient interactions or due to CCR4-NOT complex partitioning into distinct nanocompartments within stress granules, which would require experimental validations. Additionally, the direction of interaction change depends on the bait used as a probe. We observe increased proximal interactions between the CCR4-NOT complex and other stress granule proteins, PRRC2A and PRRC2B, indicating that the direction of change in bait-specific^109^.

The discovery of CCR4-NOT localization to stress granules motivated us to investigate whether stress granules contribute to post-transcriptional regulation—a long-standing controversy. Poly(A) tails lengthen during cellular stress^56-59^, yet the mechanism has remained unclear. We hypothesized that sequestration of the CCR4-NOT complex into stress granules inhibits its deadenylase activity, thereby driving poly(A) lengthening. Under basal conditions, CCR4-NOT resides in the cytoplasm, not in P-bodies, and modulates transcriptome-wide poly(A) tail lengths, but during stress, it is sequestered into stress granules, away from the translationally active cytosol, where its activity may be regulated.

To test this model, we performed global transcriptomics analysis in three conditions: no stress, oxidative stress, and oxidative stress with lipoamide (in which CCR4-NOT remains in the cytosol; Figure 6A). Blocking stress granules and releasing CCR4-NOT during stress resulted in global poly(A) tail shortening, reversing the poly(A) lengthening typically induced by stress (Figure 6B). Despite CCR4-NOT enrichment in stress granules, stress granule-associated transcripts also displayed increased poly(A) tail lengths, supporting that CCR4-NOT activity is inhibited in stress granules.

Statistical analysis revealed a subset of transcripts whose poly(A) tail lengths or abundance were significantly regulated by stress granule assembly (Figures 6D and 6F). Stress-induced changes in these transcripts were reversed when stress granule assembly was blocked. Transcripts whose poly(A) tails lengthened in a stress granule-dependent manner were enriched for genes involved in transcription and stress adaptation, including chaperones and components of the MAPK signaling pathway. We propose that CCR4-NOT complex sequestration into stress granules augments the cellular stress response by increasing the levels of key transcripts important for cellular adaptation and survival (Figure 6H).

Together, our data support a spatial mechanism by which stress granules remodel post-transcriptional regulation during cellular stress. We show that the CCR4–NOT deadenylase complex is robustly recruited to stress granules, coinciding with poly(A) lengthening and increased abundance of stress-induced survival transcripts. Disruption of stress granule assembly reverses these transcriptomic changes, indicating that condensate formation is necessary for this regulatory switch, likely through CCR4-NOT deadenylase localization. These findings suggest that stress granules function as dynamic sinks that transiently limit cytosolic deadenylation, thereby maintaining elevated levels of stress-induced transcripts available for protein production. Although direct measurements of CCR4-NOT enzymatic activity within stress granules will be required to substantiate this model, the greater extent of poly(A) tail lengthening observed in stress granule-enriched transcriptome supports the possibility that granule-localized CCR4-NOT is functionally attenuated (Figure 6B, H). More broadly, our findings support a model in which biomolecular condensates regulate gene expression not primarily as sites of biochemical activity, but by redistributing limiting regulatory factors in space and time. This work provides a mechanistic framework for understanding how condensate-mediated protein sequestration contributes to adaptive transcriptome remodeling during stress.

## LIMITATIONS OF THIS STUDY

We report dynamic proximal interaction profiles of multiple stress granule proteins utilized as molecular probes for stress granule condensate. While we are confident of proximal associations reported here, we cannot discern where these proximal interactions occur (*i.e.,* inside or outside microscopically visible stress granules). Furthermore, while we observe clear stress-context dependent heterogeneity in stress granule interaction network, we are unable to test if these interactions occur in stress granules to support structural or functional heterogeneity of stress granules. Thus, further analysis and experimentation are required to determine context-specific stress granule proteome. As with any screening approach, our proteomics and genetic screening methods contain false negatives.

With regards to the effect of poly(A) tail lengths affecting stress granule formation, we utilized cordycepin to shorten poly(A) tails globally. Poly(A) tail regulation is a dynamic process, involving both lengthening and shortening. While cordycepin shortens poly(A) tails of the transcriptome globally by terminating RNA synthesis and blocking addition of poly(A) tails ^92^, it is possible that the concentration used may affect the deadenylation kinetics, as reported in an *in vitro* study^110^, which may affect the interpretation of our experiments where we combined cordycepin and CNOT2 knockdown.

## METHODS

### Cell culture and stable cell line generation

HEK293 Flp-In T-REx (Invitrogen, R780-07; used for fixed-cell microscopy and BioID experiments), HeLa Flp-In T-REx (Arshad Desai Lab), and HeLa (Anne-Claude Gingras Lab) cells were grown at 37°C in DMEM high glucose media supplemented with 5% Fetal Bovine Serum (FBS), 5% cosmic calf serum, and 100 U/mL penicillin/streptomycin (growth media). Cells were regularly passaged upon attaining 80-100% confluency and monitored for mycoplasma contamination.

All expression constructs were created using Gateway LR cloning (Thermo Fisher). Entry clones for each bait protein were recombined with a destination vector (pDEST-3xFLAG-miniTurbo for BioID and pDEST-mCherry for microscopy-based assays (gifts from Anne-Claude Gingras, Lunenfeld-Tanenbaum Research Institute) to obtain expression constructs.

To generate stable cell lines, our expression constructs were transfected into Flp-In T-REx parental cells following a modified jetPRIME® protocol (Polyplus, CA89129-924). Briefly, cells were seeded at 25-30% confluency in a 6-well plate. The following day, cells were transfected with a mixture of 1 µg pOG44 DNA, 100 ng bait construct DNA, and 2 µL jetPRIME reagent in 200 µL jetPRIME buffer (day 2). On day 3, cells were passaged to 10 cm dishes and grown to 80% confluency, where successful transfectants were selected for using Hygromycin B-supplemented growth media (200 µg/mL; Wisent, 450-141-XL) starting on day 4. Selection media was changed every 2-3 days, until clonal colonies were visible; clonal colonies were pooled together to generate stable cell lines.

### Sample preparation for stress BioID and total proteome analysis

For BioID, we empirically tested 15-minute, 30-minute and 1 h biotin labeling using cells expressing miniT-G3BP1 and found that 30-minute labeling provided optimal balance between obtaining good labeling signal versus capturing dynamic changes. Briefly, HEK293 Flp-In T-REx stable cell lines stably, inducibly expressing BioID constructs, and negative control cell lines (miniT-GFP and HEK293 parental line) were seeded into 10 cm dishes at 30% confluency in 10 mL growth media. After 24 h, protein expression was induced with tetracycline (1 µg/mL) in biotin-depleted growth media for 20 h. Biotin-depleted media is standard growth media prepared with biotin-depleted FBS and cosmic calf serum (as reported in Schreiber *et al.,* 2024^111^). Biotin labeling was then performed for 30 mins using growth media supplemented with 50 µM biotin in (1) untreated cells, (2) stress-treated cells, and (3) stress-recovering cells. Stress, recovery, and labelling timelines follow the steps summarized in Figure S2E. For alternative NaCl stress timepoint (T2; 5-35 min), biotin labeling began 5 min into 0.2M NaCl stress for 30min following the same regime as oxidative (As(III)) stress and recovery timepoint (T3; 35-65 min), biotin labeling was performed concomitantly with fresh media change after 35 min of 0.2M NaCl stress (Figure S3B). For global proteome profiling under stress conditions, HEK293 Flp-In T-REx cell lines stable cell lines inducibly expressing miniTurbo-G3BP1 constructs were used. Cells were subjected to (1) no treatment, and (2) stress treatments following the same protocol described above. For lipoamide stress BioID, bait expression was induced for 24h in biotin-depleted media. The next day, cells were pre-treated with either DMSO (carrier) or 100 µM lipoamide for 1h prior, then exposed to 1mM or 0mM of potassium arsenate (As(V)) for 1h, after which 50 µM biotin was added for 30min of labelling. After labelling, cells were rinsed with phosphate-buffered saline (PBS) and lifted using 0.05% trypsin. Harvested cells were rinsed with PBS, transferred to pre-weighed 1.5 mL Eppendorf tubes, pelleted, and flash frozen on dry ice and stored at -80°C.

### Generation of Pooled CNOT6L/8 Knockout Cells

Single guide RNAs for CNOT6L and CNOT8 were designed using the CRISPick online tool (https://portals.broadinstitute.org/gppx/crispick/public) with the following settings: Reference Genome = Human GRCh38 (NCBI RefSeq v.GCF_000001405.40-RS_2024_08), Mechanism = CRISPRko, Enzyme = enAsCas12a. A double-stranded geneBlock comprising the CNOT6L and CNOT8 sgRNAs separated by a 20 nt direct repeat sequence and flanked by homology arms (5’-AGGACGAAACACCGGTAATTTCTACTCTTGTAGATGGTGCAGCGCTGTCAAGTGTGTCTAATTTCTACTATTGTAGATAAAGGTCAACATTGCACCGCAGAAAATTTCTACTCTAGTAGATATAGATCTCCTTGCTAACTCAGGTAATTTCTACTGTCGTAGATATTTCTATGGTGTTACCTGAAGATTTTTTGAATCTGGTCTTGAAAAAGTGGCACCGAGTCGGTGA-3’) was obtained from Integrated DNA Technologies. Three micrograms of the enAsCas12a-encoding vector pRDA-550^112^ (a gift from Dr. Diane Haakonsen, Lunenfeld-Tanenbaum Research Institute) was incubated with 10 units of the restriction enzyme Esp3I (New England Biolabs) and 1 unit of recombinant shrimp alkaline phosphatase (New England Biolabs) at 37°C for 2 h and purified by gel extraction. Knockout constructs were generated by Gibson assembly^113^ in 20 µL reactions containing 0.02 pmol of digested pRDA-550 vector, 0.06 pmol of the double-stranded sgRNA, 0.08 units of T5 Exonuclease (New England Biolabs), 0.5 units of Phusion DNA Polymerase (New England Biolabs), 80 units of Taq DNA Ligase (New England Biolabs), 5% PEG-8000, 100 mM Tris-HCl pH 7.5, 10 mM MgCl_2_, 10 mM dithiothreitol, 1 mM nicotinamide adenine dinucleotide, and 0.2 mM each of dATP, dCTP, dGTP, and dTTP, incubated at 50°C for 1 h. After transforming the reaction into chemically-competent *Escherichia coli* NEB10 cells, single clones were selected and validated by sequencing.

For knockout cell line generation, HeLa Kyoto cells stably expressing GFP-tagged FUS (FUS-GFP^114^) were transfected with a plasmid encoding either EnAsCas12a only or EnAsCas12a plus sgRNAs targeting CNOT6L and CNOT8 using the Lipofectamine 3000 kit transfection protocol (Invitrogen). Cells were selected with 2 µg/mL puromycin after 48 h. A pool of CNOT6L and CNOT8 knockout cells was used for either scrambled or combined esiRNA-mediated knockdown experiments.

### EsiRNA Knockdown of CNOT 2, 6, and 7

CNOT6L/8 knockout (KO) cells or Cas12a only cells were seeded into 12-well plates onto coverslips at a density of 15,000-20,000 cells/mL. The next day, cells were transfected with esiRNA for CNOT6, CNOT7, CNOT2, or scrambled esiRNA in freshly provided media without antibiotics using Lipofectamine 3000 reagent. Briefly, each esiRNA (125 ng) was incubated in Opti-MEM (125 μL) at room temperature for 5 min, alongside a separate mixture of Lipofectamine 3000 reagent (1.25 μL) and Opti-MEM (125 μL) per well. Then, the two mixtures were combined and incubated at room temperature for 20 min. The transfection mixture (250 μL/well) was added to cells, and the media was changed 5 h post-transfection to fresh growth media without antibiotics. The cells were transfected a second time 24 h later, in the same manner as described above. At 24 h post-2^nd^ transfection, the media was refreshed (no antibiotics). After two esiRNA transfections (48 h post 2^nd^ transfection), the cells were then treated with Lipoamide and As(V) in the same manner as described in ‘RNA Collection, Processing and Nanopore Direct RNA Sequencing’ section. The coverslips were then removed for immunofluorescence analysis, and the remaining cells from untreated samples were harvested for S-Trap sample preparation for total proteome MS analysis and extraction of protein abundance for CCR4-NOT complex components.

### Puromycin Incorporation Assay

HEK293 Flp-In T-REx cells were plated on a 6-well plate. For each time point, puromycin was incubated for 10 min at a final concentration of 10 μg/mL, as described in Schmidt *et al.*^115^. Cells were either untreated, stressed, or stressed then recovered and puromycin incorporation was performed at indicated condition and timeframes in Figure S2F. Cells were then lifted using trypsin and collected in pre-weighed centrifuge tubes. Cells were normalized by pellet weight and lysed with 2% SDS in Tris-HCl pH 7.5. Samples were boiled with Laemmli sample buffer for 5 min at 95°C, then analyzed by Western blot (Puromycin Antibody, clone 12D10, Sigma, MABE343).

### BioID bait selection process

Bait proteins were selected for BioID experimentation by optimizing for the maximum number of theoretical bait-prey interactions using published BirA* BioID data acquired under no stress^63^. We found that 58 baits profiled during no stress showed high specificity in their ability to label and identify literature-curated RNA granule proteins (*i.e.,* ≥70% of their high-confidence interactors are annotated as Tier 1^116^). Using the concepts of the greedy heuristic, we predicted that using five baits (DAZL, YTHDF2, DDX3X, CNOT2, EIF4A2) would recover ∼75% of the literature-curated Tier 1 stress granule proteome^116^. The previous study identified stress granule proteins that dually localize to stress granules and other compartments, like P-bodies and nuclear ribonucleoprotein (RNP) complex^63^. To capture components that dynamically transit between stress granules and other RNA-containing compartments, we included CELF1 and ZC3HAV1, which identify stress granule proteins and proteins in the nuclear RNP complex or P-bodies, respectively. Next, G3BP1 and FMR1 were added for their importance in stress granule dynamics and translation regulation, respectively. The final list and their relevant selection criteria are summarized in Table S2.

### BioID sample purification for mass spectrometry

Cell pellets were thawed and resuspended in ice-cold modified RIPA lysis buffer (50 mM Tris-HCl [pH 7.5], 150 mM NaCl, 1 mM EGTA, 1.5 mM MgCl_2_, 1% NP-40, 0.1% SDS) + protease inhibitor cocktail at a 1:10 milligram-to-millilitre ratio. Samples were sonicated for 30 s (10 s on, 5 s off, repeat x3) at 30% amplitude using a 1/8” microtip at 4°C. 250 U of Benzonase Nuclease was added to each sample, which was then rotated end-over-end for 15 min at 4°C. Sample SDS concentration was raised to 0.4%, then rotated end-over-end for 5 min at 4°C. Lysate volume was normalized to the lowest weight sample within a bait-stress group, then centrifuged at 20,000 x g for 20 min at 4°C. Lysates were added to 15 µL of modified RIPA-washed Streptavidin-Sepharose beads of ∼60% slurry (Cytiva, 17511301), then rotated end-over-end for 3 h at 4°C. Beads were pelleted by centrifugation at 500 x g for 1 min, the supernatant aspirated, and the beads transferred to a new 1.5 mL Eppendorf tube using 1 mL modified RIPA buffer. Next, beads were subject to consecutive 1 mL washes (1 x SDS wash (50mM Tris-HCl [pH 7.5]), 2 x modified RIPA washes, 3 x ammonium bicarbonate (ABC) washes (200 mM ABC)), pelleting and aspirating off wash solutions in between. Beads were resuspended in 1 µg trypsin in 30 µL of 50 mM ABC solution for overnight on-bead digestion at 37°C with agitation. The next day (16 h after trypsin addition), 0.5 µg trypsin in 10 µL of 50 mM ABC was added to each sample, allowing the samples to digest for a further 3.5 h at 37°C. Beads were pelleted by centrifugation at 400 x g for 2 min, the supernatants containing released peptides were collected into a final collection tube, and the beads were washed with high-performance liquid chromatography (HPLC)-grade water twice, pelleting the beads and collecting the supernatants into the final collection tubes each time. The final collection tubes, carrying the total peptide supernatants, were centrifuged at 16,100 x g for 10 mins, and the top 80 µL was transferred to new tubes, which were acidified to 2% formic acid to stop tryptic digestion and dried down in a centrifugal vacuum concentrator.

### S-Trap sample preparation for total proteome MS analysis

Protein digestion was performed using S-Trap^TM^ micro spin columns (ProtiFi) following the manufacturer’s protocol with minor modifications. Frozen cell pellets were thawed and resuspended in 1X lysis buffer (5% SDS, 50 mM TEAB, pH 8.5) at a ratio of 100 μL buffer per 10 mg pellet. After sonication, samples were centrifuged at 13,000 x g for 8 min, then the supernatant was transferred to a new tube and the protein concentration was measured with a Pierce^TM^ BCA Protein Assay Kit (Thermo Fisher Scientific). For each sample, 50 µg of lysate protein was reduced with 20mM dithiothreitol at 95°C for 10 min, then alkylated with 100 mM iodoacetamide at room temperature for 30 min in the dark. Samples were then acidified with phosphoric acid to pH≤1. Subsequently, 505 μL of binding/wash buffer (100mM TEAB in 90% methanol) was added to the solution. This mixture was loaded onto S-Trap micro spin columns and centrifuged at 4,000 × g for 30 s to trap proteins. Following the first centrifugation, washing was repeated four times by adding 150 μL of binding/wash buffer and centrifuged at 4000 × g for 30 s. The final column spin was conducted at 4000 × g for 1 min to fully remove binding/wash buffer.

Next, 20 μL of trypsin (∼1:10 enzyme/protein) in digestion buffer (50 mM TEAB) was added to the surface of the filter and incubated for 2 h at 47°C. The tryptic peptides were eluted sequentially with 40 µL each of (1) 50 mM TEAB, (2) 0.2% formic acid, and (3) 50% acetonitrile via centrifugation for 1 min at 4000 × g. Eluted solutions were pooled together and the final peptide concentration was measured with a colorimetric Pierce^TM^ Quantitative Peptide Assay Kit (Thermo Fisher Scientific). Eluates were dried down via vacuum centrifugation and stored at −80°C.

### Liquid Chromatography-Mass Spectrometry

BioID peptides were resuspended in 12 µL of 5% formic acid and centrifuged at >20,000 x g for 1 min. For MS analysis of BioID peptides, each sample was further normalized to the smallest pellet weight across all bait-stress groups by diluting ∼1-2 µL of the peptide solution with 5% formic acid to a volume of 12 µL. Subsequently, 1 µL of 50 fmol/mL Pierce Retention Time Calibration (iRT) peptides was added, bringing the final volume to 13 µL. For MS analysis of total proteome samples, peptides were resuspended in 5% formic acid to 200 ng/µL, and 1 µg of peptide suspension was combined with 1 µL of 50 fmol/mL Pierce iRT peptides in a total volume of 13 µL.

For each sample, 6 µL of peptides was loaded onto a 10.5 cm 75 µm ID emitter tip packed in-house with 3 µm ReproSil Gold 120 C18 (Dr. Maisch HPLC GmbH) using an Easy-nLC 1200 liquid chromatography system. Peptides were eluted at 200 nL/min over a 120 min gradient, starting with 2.4% acetonitrile in 0.1% formic acid and gradually increasing to 35% acetonitrile. The acetonitrile concentration was further increased to 80% over an 8 min gradient and maintained at this concentration for 16 min at 200 nL/min, for a total run time of 144 min. Samples were analyzed with an Orbitrap Exploris 480 (Thermo Fisher Scientific) in both Data-Independent Acquisition and Data-Dependent Acquisition modes. Instrument configurations for both acquisition modes were set up using Thermo Xcalibur (V4.6.67.17) software. The instrument was operated with a positive polarity ion voltage of 2.4 kV and the ion transfer tube temperature was set to 300°C. MS analyses of each BioID sample was done in both DIA and DDA modes consecutively, while the fractionated HEK293 peptides were analyzed in DDA mode only.

For DDA experiments, MS1 scans ranged from 390-1010 m/z were performed at a resolution of 120,000, with a 3-second cycle time. The normalized automatic gain control (AGC) target was set to 125%. Precursor ions with charges from 2+ to 7+ that exceeded an intensity threshold of 1.0 × 10³ were selected for fragmentation with dynamic exclusion enabled for 9 seconds. Fragmentation was achieved with a normalized collision energy (NCE) of 33%. MS2 scans were performed at a resolution of 15,000, with a normalized AGC target of 400%, and a maximum injection time of 50 ms. The DDA-MS data were used to generate spectral library for DIA deconvolution and benchmark BioID-DIA analysis workflow.

For DIA experiments, a staggered window scheme was utilized as described in Pino et al., 2020^117^. Briefly, this approach employed two alternating precursor isolation patterns with the m/z ranges of 400-1,000 and 396-1,004 m/z, respectively. MS1 scans were acquired in centroid mode at a resolution of 60,000, a normalized AGC target of 100%, a maximum injection time of 55 ms and a scan range of 390-1,010 m/z. MS2 scans were acquired in a DIA window scheme consisting of 75 and 76 8m/z isolation windows, at a 15,000 resolution, with a scan range of 145-1450 m/z, a normalized AGCw target of 1000%, a maximum injection time of 23 ms and normalized collision energy of 33%.

The Gas Phase Fractionation (GPF) approach was chosen over the DDA-based library generation for a subset of experiments because it reduced the time and effort required. To generate a spectral library for the analysis of global proteome changes under stress, six GPF acquisitions were acquired from a pooled biological sample pool from HEK293 cells (2 µL each from 15 total proteome samples). The GPF acquisition scheme utilized was described in detail in Pino et al., 2020^117^. Each GPF injection was acquired at a MS1 resolution of 60,000 and MS2 resolution of 30,000, with an AGC target of 1000%, maximum injection time of 55 ms, and normalized collision energy of 33%. Precursor ions were isolated using staggered 4 m/z windows. Each of the six injections covers a distinct m/z range: 398–502, 498–602, 598–702, 698–802, 798–902, and 898–1002.

### Mass Spectrometry Data Analysis: Data-Dependent Acquisition and Gas Phase Fractionation

All MS files acquired in DDA and GPF modes were analyzed separately with FragPipe (v20.0), MSFragger(v3.8)^118^, IonQuant (v1.9.8)^119^ and Philosopher (v5.0.0)^120^. For GPF data, Thermo RAW files were first computationally demultiplexed and converted to mzML format prior to analysis (see **Demultiplexing of DIA staggered windows**). The protein sequence database contains reviewed UniProt entries and 15 iRT peptides as well as the same number of reversed decoy sequences, with a total search space of 40,884 sequences. The precursor mass window was set at ±20ppm and the fragment mass tolerance was set at ±20ppm. Tryptic digestion was specified with full enzyme specificity, allowing up to two missed cleavages. Methionine oxidation (+15.9949 Da), and N-terminal acetylation (+42.0106 Da) were set as variable modifications, while carbamidomethylation of cysteine (+57.0215 Da) was set as a fixed modification. For BioID samples, carbamidomethylation of cysteine (+57.0215 Da) fixed modification was omitted, as alkylation step is not included in BioID processing steps. Additionally, peptide N-terminus heavy arginine (+10.008269) and heavy lysine (+8.014199) were included as variable modifications to facilitate the identification of iRT peptides. The GPF data search also included lysine biotinylation (+226.07760) as a variable modification. MSBooster was enabled, which uses deep learning to predict retention time and spectra to increase identifications^121^. PSM validation was done using Percolator (--only-psm --no-terminate --post-processing-tdc), which uses a linear support vector to performs FDR calculations using a target-decoy approach^122^. This is followed by protein inference by ProteinProphet (--maxppmdiff 20)^123^. FDR filtering was done using Philosopher to 1% PSM and protein levels (--sequential --picked --prot 0.01).

MS1 intensity-based quantification was conducted using IonQuant with Match-Between-Runs (MBR) enabled (default parameters)^119,124^. MaxLFQ intensity (with minimum ions set to 2) was used for downstream statistical analysis. For feature detection and peak tracing, m/z tolerance was set to 10ppm and RT Window to 0.6. MBR top runs was set to 10.

### Spectral Library Generation

Two spectral libraries were generated for DIA data analysis: (1) DDA-based library used for the analysis of BioID data and (2) GPF-based library for global proteome analysis. For the first library, the DDA search results from fractionated peptides of HEK293 lysate and BioID peptides were compiled into a spectral library using EasyPQP with default parameters (v0.1.42) (https://github.com/grosenberger/easypqp). The initial library file then underwent a 3-step filtering process to generate the final library in TSV format. First, precursors containing alkylated cysteine residues were filtered out to minimize false positives because cysteine reduction and alkylation is not performed during BioID sample processing. Second, fragments outside the 350-2000 m/z range were removed as signals outside this range are typically noisy. Finally, the six most intense fragments for each precursor were retained, and precursors with less than six fragments were excluded^125^. Using a defined number of fragments in the library ensures uniform statistical distributions during automated peak scoring in DIA analysis (i.e., different numbers of fragments per precursor may result in mixed statistical distributions for the target identifications). The final DDA-based spectral library contains 193,510 precursors from 9,860 proteins.

The generation of GPF-based library used the same workflow applied to the GPF search results. In this case, alkylated cysteine filtering was omitted, as the total proteome samples were reduced with iodoacetamide. The GPF-based spectral library contains 44,887 precursors from 37,204 peptides and 5,203 proteins.

### Demultiplexing of DIA staggered windows

DIA and GPF-MS data acquired using the staggered window scheme require a computational demultiplexing step before they can be processed by data analysis software. This step resolves the staggered precursor isolation windows into effective windows that are halves of the original width. For example, 8 m/z staggered windows spanning m/z range of 400-100m/z would be demultiplexed into non-overlapping 4m/z windows: 400–404 m/z, 404–408 m/z, 408–412 m/z, etc. The demultiplexing was performed by MSConvert (ProteoWizard) with the following filters: 1) ”peakpicking vendor msLevel=1-“; 2) “demultiplex optimization=overlap_only massError=10.0ppm”. The resulting mzML files are used for downstream data analysis

### Mass Spectrometry Data Analysis: Data-Independent Acquisition

The DIA-MS data in mzML format were analyzed by the DIA-NN software (v1.8.1)^81^ using a library-based approach. BioID and global proteome data were analyzed separately. The BioID and CNOT KD total proteome data were searched against the DDA-based spectral library described above (see **Spectral Library Generation**), while analysis of the stress total proteome data utilized the GPF-based spectral library. DIA-NN was configured with match-between-runs enabled, cross-run normalization and MaxLFQ-based protein quantification disabled. Mass accuracy was set to 20 ppm for both MS1 and MS2. Additionally, the *“--report-lib-info*” flag was set to generate fragment-level intensity values in the resulting report.tsv file, which was used for downstream statistical analysis. The resulting DIA-NN report.tsv file was first filtered using global precursor and protein FDR thresholds of 1% (Lib.Q.Value ≤ 1% and Lib.PG.Q.Value ≤ 1%) to ensure high-confidence identifications. Fragment-level (MS2) intensity data were extracted from the “Fragment.Quant.Raw” column in the report.tsv.

For BioID data, intensity normalization was then performed within each bait group by scaling fragment intensities to a common total MS2 intensity. Specifically, the intensity value of each fragment in a sample is divided by the sum of total fragment intensities in the sample and multiplied by the median of the sum of fragment intensities across all samples in a bait or control group. The resulting normalized fragment intensity matrix was used for SAINTq (V0.0.4)^82^ analysis to identify high-confidence interactors and for Mixed-Model ANOVA to identify dynamically associating interactors (see Data Analysis). For global proteome data, intensity normalization was performed by adjusting for total MS2 intensity across all samples prior to mixed-model ANOVA to identify differentially abundant proteins.

### Mass Spectrometry Data Deposition

Datasets consisting of raw files and associated peak list and results files have been deposited in ProteomeXchange through partner MassIVE as complete (Data Dependent Acquisition) or partial (Data Independent Acquisition) submissions. Additional files include the complete SAINTexpress (V3.6.3) outputs for each dataset and a “README” file that describes the dataset composition and the experimental procedures associated with each submission. The DDA and DIA datasets generated here were each submitted as independent entries. The accession numbers for the DDA dataset are MSV000095953 (ftp://massive.ucsd.edu/v08/MSV000095953/) in MassIVE and PXD056247 in ProteomeXchange. The accession number for the DIA dataset is MSV000095944 (ftp://massive.ucsd.edu/v08/MSV000095944/). The accession number for the total proteome DIA dataset including GFP-DIA data used to generate spectral library is MSV000098662 (ftp://massive-ftp.ucsd.edu/v10/MSV000098662/). Total proteome data for cell lines with knockouts and/or knockdowns of CCR4-NOT components were deposited with the MassIVE ID MSV000100193 (ftp://massive-ftp.ucsd.edu/v11/MSV000100193/).

### Data Analysis: SAINTexpress and SAINTq to Identify Confident Interactors

Significance Analysis of INTeractome (SAINT) comprises a series of computational tools used to assign confidence scores to protein-protein interaction based on quantitative data generated from AP-MS and BioID experiments^65^. Recent versions of SAINT have been extended to analyze intensity-based data^66,82,126^, with SAINTq implemented specifically for the analysis of intensity data at the protein, peptide and fragment levels^82^. Although SAINTq has been available since 2015, its application in BioID projects has not been widely tested. We generated BioID-SAINTq analysis pipeline for protein- and fragment-level intensity data and benchmarked it against the established BioID-SAINTexpress.

#### SAINTexpress Workflow

Spectral-count based analysis of DDA BioID data was performing using SAINTexpress as implemented within the Fragpipe framework. The “reprint.spc.tsv” was the input for SAINTexpress and the analysis was performed using the default parameters. Identified prey proteins with BFDR ≤ 0.05 are considered high-confidence interactors.

#### SAINTq Workflow

For DDA protein-level intensity-based analysis used for internal quality control, we used the “combined_protein.tsv” file generated by Fragpipe. After selecting the MaxLFQ intensity columns, the data were normalized by total intensity within each bait group before running a modified version SAINTq (v0.0.4), provided by the Anne-Claude Gingras Lab (https://github.com/vesalkasmaeifar/Modified_SAINTq_and_SAINTexpress). For DIA fragment-level intensity-based analysis, the total intensity normalized fragment matrix (see Mass Spectrometry Data Analysis: Data-Independent Acquisition) was analyzed by SAINTq. In both DDA and DIA SAINTq analyses, control compression was disabled and default parameters were applied. Using the benchmark of known stress granule proteome and results from DDA SAINTexpress, we have manually assigned BFDR 0.03 as a suitable threshold for high-confidence interaction.

### Data Analysis: Unsupervised Hierarchical Clustering and Data Visualization

To ensure the quality of DIA proteomics data prior to unsupervised hierarchical clustering, we applied a two-step filtering process to each bait group. First, fragments missing in more than 80% of samples within a group were excluded from the analysis. Second, precursors with fewer than three fragments were filtered out. These filters were applied because fragments or precursors not quantified in the majority of samples are likely noises. Next, protein intensity values were rolled up by summing intensities of corresponding fragments and the resulting intensity values were log2-transformed.

To infer proximal interactions associated with the stress granule condensate rather than bait-specific interactions, we filtered for 1,801 high-confidence interactors (BFDR ≤ 3%, SAINTq) that were identified by at least two baits, regardless of the condition. We assumed that high-confidence interactors identified across multiple baits are more likely to be associated with stress granules.

Clusterings of bait-conditions (columns) and proteins (rows) was performed using Euclidean distance and Ward’s method for linkage. The R package pheatmap was utilized for heatmap visualization. To determine the most informative number of clusters for protein groupings, we computed the within-cluster sum of squares (WCSS) across a range of cluster numbers. WCSS measures the total variance within each cluster by summing the squared distances between each protein and the average profile of its assigned cluster. A lower WCSS value indicates more compact and coherent groupings, reflecting similar abundance patterns across samples. An elbow plot was generated to visualize the change in total WCSS as the number of clusters increased. The optimal number of clusters was selected at the point where the curve plateaued. Based on this approach, four clusters were identified as the most informative. Finally, gene ontology (GO) enrichment analyses were performed for proteins in the four-row clusters using g:Profiler. The background protein set for the GO enrichment analysis consisted of 9,860 protein entries in the spectral library, as these represent the proteins detectable in the experiments. GO terms with FDR ≤ 1% (Benjamini-Hochberg) are significant.

### Data analysis: Mixed-Model Analysis of Variance

The total intensity normalized fragment matrix contained a sizable proportion of missing values. To improve statistical power, the matrix was imputed using a K-Nearest Neighbor (KNN) approach using the python package scipy (V1.10.1), with the number of neighbors (k) set to 5. We then used the filtered proteins from two-step filtering process (see Data Analysis: Unsupervised Hierarchical Clustering and Data Visualization), to select for high-quality fragments. Next, precursor intensity values were rolled up by summing intensities of corresponding fragments and the resulting intensity values were log2-transformed.

A linear mixed model was then applied to the KNN-imputed, log₂-transformed precursor intensity matrix. This model was used separately to the BioID and global proteome datasets to assess prey protein dynamics across time points for each bait (**T1 vs T2_As(III) vs T3_As and T1 vs T2_NaCl vs T3_NaCl**), and to evaluate global proteome changes in response to stress (**T1 vs T2_As(III) vs T2_NaCl**), respectively.

The linear mixed model is defined as:

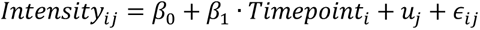

where *Intensity*_*ij*_ represent log2-transformed intensity for precursor *j* at time point *i*. *β*_0_ is the overall intercept, *β*_1_is the fixed effect coefficient representing the effect of time point *i*, *u*_*j*_ is the random effect accounting for variability among different precursors, and *ϵ*_*ij*_ is the residual error term. This model accounts for the random effect of the precursors corresponding to the same protein and the fixed effect of the timepoint. In cases where only a single precursor is available, a simpler linear model is applied without the random effect:

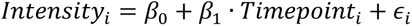

Analysis of variance (ANOVA) was then applied to obtain p-values indicating the significance of time point effects. Post-hoc analysis was conducted using estimated marginal means to evaluate pairwise fold change between time points (i.e T1 vs T2_As(III)). The Benjamini-Hochberg procedure was applied to correct for multiple comparisons, with differential interactions defined by p-adjusted values ≤ 0.01 and fold change ≥ 1.5.

### Immunofluorescence (IF) microscopy

For bait localization and prey colocalization tests, stable HEK293 or HeLa Flp-In T-REx BioID cell lines were seeded onto glass coverslips in 6-well plates at 20% confluency in 2 mL growth media, and protein expression was induced 24 h later with tetracycline (1 µg/mL) for 20 h. For lipoamide titration tests, HEK293 BioID stable cells stably expressing miniT-G3BP1 or miniT-GFP were seeded at 30% confluency, then induced for bait expression for 24 h. Varying concentrations of lipoamide (0-100 µM) or carrier (DMSO) were added, followed by no treatment (untreated) or potassium arsenate (As(V)) stress (1 – 2 mM). For stress granule assembly assays in cordycepin treatment (Figure 4A-B), HEK293 Flp-In T-REx, HeLa Flp-In T-REx, or HeLa cells were seeded onto glass coverslips in growth media. Glass coverslips were treated with poly-D-lysine for 1 h at 37°C prior to seeding HEK293 Flp-In T-REx cells. Cells reached 90% confluency 24-48 h post seeding onto the glass coverslips and were treated with either a media change (untreated), sodium arsenite (III) [60 µM – 1 h, 75 µM – 1 h, 500 µM – 30 min], sodium chloride [0.2 M – 1 h], or sorbitol [sorb: 0.4 M, 1 h]) at 37°C.

To detect colocalization of 60S subunits (RPL32 and RPL36) with stress granules in Figure S7, HeLa cells were treated with either sodium arsenite (III) (0.5mM) or sodium chloride (0.2M) for 30 min, washed once with ice-cold PBS, then incubated twice with 0.01% digitonin in PBS on ice for 2 min. After the second incubation, the digitonin was aspirated, and the cells were fixed and processed for IF processing steps below. For the cordycepin treatment in Figures 4A-B and S8C-D, cells were pretreated with varying concentrations of cordycepin [0-100 µM; C3394, Sigma-Aldrich] for 3 h, with varying amounts of sodium arsenite (III) added during the last h before fixation. For the cordycepin treatment in Figure 4D, the HeLa Kyoto FUS-GFP cells (MCB_005340) were first treated with the indicated concentration of cordycepin (C2689, Tokyo Chemical Industry) for 1 h, then treated with 10 µM of lipoamide or the corresponding percentage (v/v) of DMSO (0.1%) as the control for 1 h, and finally stressed with 1 mM of potassium arsenate (As(V)) for 1 h before fixation with 4% formaldehyde in PBS.

Cells were processed for IF immediately following drug treatments by rinsing the cells with PBS twice, then fixing with 4% paraformaldehyde in PBS for 10-15 min. Post fixation, cells were washed 3 times with PBS or TBS-T and then permeabilized with -20°C methanol (Figure 4A, 5, S2B, S7A-B, S8, S9A-B, S10, S11A) or 0.1% Triton-X in TBS-T (Figure S1B, S2A, S2C, S2D, S6C, S9C-D) at room temperature for 10 min. Cells were then washed 3 times with PBS or TBS-T, then blocked with 5% milk in TBS-T or 5% BSA in PBS for 1 h. Coverslips were then incubated in primary antibody solution prepared in 5% BSA in PBS for 1 h or overnight (depending on the antibody) in a humidity chamber at 4°C. The cells were then washed 3 times with PBS or TBS-T and then incubated with secondary antibodies and DAPI prepared in 5% BSA or milk blocking solution for 1 h in the dark. Coverslips were then washed for 3 times with PBS or TBS-T. Finally, samples were mounted onto slides using Prolong Gold and cured in the dark overnight. All steps were performed at room temperature, excluding the overnight incubation. Samples were imaged using the Leica SP8 Confocal Microscope and the Leica Stellaris 5 Confocal Microscope at 40X and/or 63X magnification using LASX (V3.5.7.23225 and 4.7.0.28176), and images were processed using either IMARIS (V10.1.1) or LASX software (4.8.0.28989).

### High Content Genetic Screen of Lipoamide

To generate a cell line stably expressing Cas9 and GFP-tagged FUS, A plasmid of human codon-optimized *S. pyogenes* Cas9 (lentiCas9-Blast; #52962, Addgene) was transfected into HeLa Kyoto cells stably expressing FUS-GFP^114^ (MCB_005340), and the clone was selected with 20 µg/mL of blasticidin.

The obtained clone cells were seeded at 550 cells/well in 384-well plates (#781092, Greiner), transfected with 20 nM each of tracrRNA (U-002005, Horizon) and sgRNA from Human Edit-R crRNA library (18,710 genes tested; Dharmacon) using Interferin (polyplus), and incubated for two days before the medium was replaced with fresh one for 1 day. CRISPR-Cas9-mediated gene knockout was validated with KIF11 sgRNA, which kills cells. An sgRNA targeting SRSF1 (CM-018672-01, Horizon) was used to validate the assay. Mock transfection was used for controls both positive and negative for stress granule assembly. Three days after transfection, the cells were pre-treated with 20 µM of lipoamide for 1 h and then stressed with 1 mM of potassium arsenate (As(V), A6631, Sigma Aldrich) for additional 1 h in the presence of lipoamide before fixed with 4% formaldehyde in PBS. For only mock transfection, wells without lipoamide treatment were also prepared for the positive control. The fixed cells were stained with 1 µg/mL of DAPI and 0.25 µg/mL of CellMask blue (H32720, Thermo Fisher Scientific) and imaged on a CellVoyager CV7000 (R1.17.05) automated spinning disc confocal microscope (Yokogawa) with a 40× NA 1.3 water objective lens.

For the secondary screen, the HeLa Kyoto FUS-GFP cells were seeded at 1000 cells/well in 96-well plates (#655090, Greiner), transfected with 25 ng of each endoribonuclease-prepared small interfering RNA (esiRNA; Eupheria) using Lipofectamine2000 (Invitrogen). Three days after transfection, the cells were treated with 10 µM of lipoamide for 1 h and then stressed with 1 mM of potassium arsenate (As(V)) in the presence of lipoamide for 1 h before fixed with 4% formaldehyde in PBS. The cells were stained with 1 µg/mL of DAPI and imaged similarly to the primary screen.

### Confocal Image Analysis

For high content imaging analysis (relevant to Figures 3B-E and 4C-D), segmentation of stress granules were performed based on cytoplasmic FUS-GFP signals using CellProfiler (V4.2.4)^127^, and the obtained information (*e.g.*, number and size of stress granules, intensity of FUS-GFP) was processed with KNIME (V4.7.0) and Fiji (V2.16.0). For primary genome-wide CRISPR screen, the number of stress granules per cell treated with each sgRNA was normalized relative to two datasets: that of mock transfection without lipoamide treatment (defined as 100%) and that of mock transfection with lipoamide treatment (0%) within each plate; the normalized values were transformed into z-scores as shown in Figure 3. For the quantification of immunofluorescence images (relevant to Figure 4A-B), segmentation and quantification were performed using CellProfiler (V3.0). Manual quantification of immunofluorescence images shown in Figure 3F was done using Fiji (V2.14.0). All of the images were processed together to avoid bias during quantification. Any cells with ≥3 FUS-GFP positive puncta in the cytoplasm were counted as a stress granule-positive cell.

For colocalization analysis between large ribosomal subunits (RPL32 and RPL36) in Figure S7, CellProfiler Version 4.2.8 was used. First, the nucleus was segmented with the ‘IdentifyPrimaryObjects’ using DAPI. Then, the nucleus was masked in the RPL channel to exclude the nuclear signal from analysis. Colocalization measurements across the entire image were performed for the masked RPL channel and G3BP1 channel using the MeasureColocalization module.

### Cyclohexamide Treatment

HeLa Flp-In cells were seeded onto coverslips in a 12-well plate with 1.5 X10^5^ cells. The following day, the cells were stressed with sodium arsenite (III) at a concentration of 60 µM. After 1 h of stress, either ethanol (carrier) or cyclohexamide (10 µg/mL final concentration) was added via spike-in for both 30 and 60 min. Cells were then processed following the IF protocol listed above.

### RNA Collection, Processing and Nanopore Direct RNA Sequencing

The following steps were applied to three biological replicates. HeLa Flp-In cells were seeded into 6cm dishes with 1X10^6^ cells, including and matching samples at the same relative density onto coverslips in a 12-well plate for IF. The next day, the cells received a fresh media change 1 h prior to adding lipoamide. Lipoamide (in DMSO) was added to the cells via media change at a concentration of 200 µM. DMSO was added as a carrier to the no-stress and sodium arsenate (As(V)) conditions. After 1 h of lipoamide, cells were stressed with arsenate (As(V) in water) via spike-in at a final concentration of 20mM. One hour after As(V) was added, the cells were collected simultaneously for IF and RNA extraction. The IF was processed as described above. The samples for RNA were washed quickly with PBS twice, then lysed with 600 µL Tri Reagent per 6 cm dish for 5 min with rocking. Samples were moved to a Lo-Bind microfuge tube on ice, and 120 µL chloroform was added to the samples. Samples were shaken vigorously 20 times and incubated on ice for 15 min. Samples were then centrifuged at 4°C for 15 min at 12,000 rpm, and the aqueous (top) layer (∼300 µL) was carefully removed and placed into a new Lo-Bind tube on ice. Following this, 300 µL of isopropanol was added to the samples, and they were shaken vigorously 20 times and incubated overnight at -20°C. The next day samples were spun at 4°C for 15 min at 12,000 rpm. The isopropanol was carefully removed, leaving a small pellet. The pellets were washed with 1 mL of ice-cold 100% ethanol and vigorously shaken 20 times. The samples were spun again at 4°C for 10 min at 12,000 rpm. The ethanol was carefully removed, and the pellet was left to dry at room temperature. Once dry the RNA pellet was resuspended with 50 µL of nuclease-free water and concentration was checked with a Biotek Epoch reader.

To prepare the RNA samples for subsequent analysis, 10 µg of RNA was digested with TURBO DNase following the manufacturer’s protocol (30 min at 37°C). The RNA samples were then cleaned up with the Qiagen MinElute Column Clean-up Kit (Cat. 74204) following the manufacturer’s protocol for Nanopore Direct RNA Sequencing or RT-qPCR.

Direct RNA sequencing was performed by The Centre for Applied Genomics at The Hospital for Sick Children, Toronto, Canada. RNA concentration and integrity were assessed using the Qubit™ RNA HS Assay Kit (ThermoFisher, Q32852) and by the Bioanalyzer RNA 6000 Nano assay (Agilent, 5067-1511). Library preparation was performed using the Direct RNA Sequencing Kit (Oxford Nanopore Technologies, SQK-RNA004) following the manufacturer’s protocol version 9195_v4_revB_20Sep2023. Briefly, 1 µg of total RNA in 8 µl nuclease-free water was used as input. Prior to reverse transcription, 25ng of *Saccharomyces cerevisiae* RNA for ENO2 was spiked into the RNA samples used for sequencing validation of ONT transcripts. Reverse transcription adapter (RTA) was ligated to the RNA using T4 DNA Ligase (NEB, M0202M) and NEBNext® Quick Ligation Reaction Buffer (NEB, B6058), followed by reverse transcription using SuperScript™ III Reverse Transcriptase (Thermo Fisher Scientific, 18080044) to synthesize a complementary cDNA strand for stability. The cDNA strand was not sequenced. Post-reverse transcription, the RNA-cDNA hybrid was purified using Agencourt RNAClean XP beads (Beckman Coulter, A63987). Sequencing adapters were then ligated to the hybrid using T4 DNA Ligase and RNA Ligation Adapter (RLA), followed by another bead-based purification step. The final library was quantified using the Qubit™ dsDNA HS Assay Kit (ThermoFisher, Q32851), and the entire volume was used for sequencing. The final libraries were sequenced on the Promethion-24 instrument (Oxford Nanopore Technologies) on FLO-PRO004RA flow cells for a 90-96 h run. Basecalling was performed with the High-accuracy model (HAC) using MinKNOW version 25.05.14 and Dorado version 7.9.8.

### Nanopore Sequencing Analysis of mRNA Poly(A) Tails and Abundance

Nanopore pod5 files were simultaneously basecalled and aligned to the hg38^128^ human reference genome using the Dorado basecaller (v0.9.0; https://github.com/nanoporetech/dorado). Poly(A) tail measurements for each read were also called with Dorado (full command: dorado basecaller hac "$pod5_files" --device cuda:0 --estimate-poly-a --reference "$reference" --mm2-opts "-k 15 -w 10" --min-qscore 9 > "$output_file"). Non-primary alignments were filtered out using samtools^129^ (v1.21). Reads were then assigned to genetic features using featureCounts^130^ (v2.1.1) and the NCBI Refseq hg38 gtf annotation file^128^. Poly(A) tail lengths were averaged across reads for each gene. Genes for which there were fewer than 10 reads were excluded from the analysis. Only reads mapping to the nuclear genome were included. Classifications for RNAs enriched in stress granules were taken from Khong et al., 2017^30^.

Differential expression analysis was conducted using DESeq2^103^. Input count tables were generated from featureCounts^129^. Scripts for this analysis can be accessed on Github (https://github.com/wormbro1/Tsai_et_al_CCR4NOT_DRS_analysis). Poly(A) length analysis was conducted using TAILCaller^101^ to identify significant changes with p_adj_ <0.05 and >|1.3|-fold in median lengths. Subsequent post-hoc condition-condition comparisons were completed based on initial test determined by TAILCaller; Dunn’s test for Kruskal-Wallis, Tukey’s HSD test for ANOVA, and Games-Howell test for Welch’s ANOVA. All post-hoc tests were corrected with Bonferroni adjustment, save for Games-Howell, which has adjustments built-in. Fold-change of poly(A) lengths was calculated from the median poly(A) length for each gene, as calculated by TAILCaller.

### RT-qPCR

100ng to 1 µg of RNA was used to prepare cDNA using qSCRIPT cDNA Synthesis Kit (QuantaBio, #95047-100) following the manufacturer’s protocol. Prior to reverse transcription, 25ng of *Saccharomyces cerevisiae* RNA for ENO2 was spiked into the RNA samples used as an internal control. RT-qPCR master mixes were prepared using the appropriate forward and reverse primers (Table S10) and PerfeCTa SYBR Green FastMix (QuantaBio, #95072-012). The qPCR was completed using a Bio-Rad CFX Connect. For all RT-qPCR experiments, 18S rRNA was used as an internal control to normalize Cq values.

## DATA AVAILABILITY

All proteomics data are available at the Mass Spectrometry Interactive Virtual Environment (MassIVE, http://massive.ucsd.edu). MassIVE ID: MSV000095953 (ftp://massive.ucsd.edu/v08/MSV000095953/); MassIVE ID: MSV000095944 (ftp://massive.ucsd.edu/v08/MSV000095944/); MassIVE ID: MSV000098662 (ftp://massive-ftp.ucsd.edu/v10/MSV000098662/); and MassIVE ID: MSV000100193 (ftp://massive-ftp.ucsd.edu/v11/MSV000100193/).

Nanopore data are available at the European Nucleotide Archive. Nanopore ENA accession number: PRJEB106356 (https://www.ebi.ac.uk/ena/browser/view/PRJEB106356).

## CODE AVAILABILITY

Data analysis and visualization codes will be shared upon request to the lead contact, Ji-Young Youn.

## ACKNOWLEDGEMENT

We thank A.C. Gingras for supervising V. Kasmaeifar; B. Larsen, C. Wong, G. Liu, and H.W. Choi for help with proteomics and SAINTq analysis; D. Haakonsen for the technical guide for using the Cas12a system; K. Talipova and L. Pelletier for their contribution to validation. We thank Tara Paton and Sachin at TCAG for their Nanopore DRS processing and thank J.L. Rubinstein for his review of the manuscript. This work was supported by the Natural Sciences and Engineering Research Council of Canada (NSERC; RGPIN-2022-04849), Canada Foundation for Innovation (CFI; 41430) and the Canadian Institutes of Health Research (481147) to J.-Y.Y, National Institutes of Health (R35GM128680) to O.R. Confocal imaging was performed at the SickKids Imaging Facility, a facility supported by Canada Foundation for Innovation, Ontario Research Fund and SickKids Research Institute. H.R. is the Canada Research Chair (Tier 2) in Mass Spectrometry-based Personalized Medicine and J.-Y.Y is the Canada Research Chair (Tier 2) in Membraneless Organelle Proteomics.

## AUTHOR CONTRIBUTIONS

Conceptualization, J.-Y.Y., A. H.; Methodology, Y.-C.T., H.U., S.R.M., E.K., K.J.S., R.B., C.M., A.J.; Software, Y.-C.T., V.K.; Validation, E.K., A.M.; Formal Analysis, Y.-C.T., H.U., S.R.M., K.J.S., J.P.T.O., J.Q.H.; Investigation, Y.-C.T., H.U., S.R.M., E.K., K.J.S., A.M., Z.Z., J.S.; Resources, C.A.L., J.Q.H.; Data Curation, Y.-C.T.; Writing – Original Draft, J.-Y.Y., H.U., Y.-C.T.; Writing – Review & Editing, H.U., Y.-C.T., E.K., A.M., K.J.S., J.Q.H., H.R., A.H., J.-Y.Y.; Visualization, Y.-C.T., H.U., J.-Y.Y.; Supervision, J.-Y.Y., A. H. H.R., O.S.R.; Project Administration, J.-Y.Y. H.R., H.U., A. H.; and Funding Acquisition, J.-Y.Y., H.R., A. H.

## DECLARATION OF INTERESTS

A.A.H. is a founder and scientific adviser board member of Dewpoint Therapeutics.

## DECLARATION OF GENERATIVE AI AND AI-ASSISTED TECHNOLOGIES

ChatGPT was employed to improve writing style and grammar checks. The authors carefully reviewed, edited, and revised the content according to their own preferences, assuming ultimate responsibility for the publication’s content.

## SUPPLEMENTAL INFORMATION

**Figure S1. Stress BioID quality control** (**a**) Western blot analysis comparing the levels of stably expressed miniTurbo-tagged proteins to the corresponding endogenous proteins. Lysates prepared from HEK293 Flp-In T-REx parental cell line, tetracycline-induced miniTurbo stable cells or uninduced cells were analyzed for each stress granule bait protein (top). Ponceau staining of the total blot is used as a loading control (bottom). Left: G3BP1 and Right: FMR1. (**b**) Immunofluorescence microscopy of miniTurbo-3xFLAG-G3BP1 cells under no stress or stressed with sodium arsenite, As(III), then BioID labeled. Stress granule marker (TIA1), bait (FLAG) and biotinylated proteins (Strep). Scale bar = 10 µM. (**c**) Lipoamide titration test to determine effective concentration for stress granule inhibition in BioID cells. Quantifications from 2,600-4,300 cells. (**d**) Condition-condition plot of miniT-G3BP1 high-confidence proximal interactors in cells incubated with a carrier (DMSO) or 100 µM lipoamide for 1 h, then biotin labelling in ‘no stress’ state. (**e**) Venn diagram comparison of high-confidence preys identified from 58 high-specificity baits from Youn *et al.,* 2018 (red), high-confidence preys of the selected nine baits (green) and literature-curated stress granule proteome (Tier 1; Youn *et al.,* 2019).

**Figure S2. Stress granule assembly and disassembly kinetics.** (**a**) Immunofluorescence microscopy images of HEK293 Flp-In T-REx cells fixed and stained for stress granules (G3BP1) at 1 h (left) or 1.5 h after recovery post oxidative stress. Scale bar = 10 µM. (**b**) Immunofluorescence microscopy images show stress granule assembly kinetics after hyperosmotic stress (0.2 M NaCl) for indicated times in HEK293 Flp-In T-REx cells. Stress granule marker G3BP1 (green); P-body marker EDC3 (red); Nuclei (DAPI). Scale bar = 10 µM. (**c**) Subcellular localization of BioID bait (FLAG), biotinylated proximal proteins (Strep), and stress granules (TIA1) after 30 min of biotin labeling in hyperosmotic stress condition in miniT-3xFLAG-G3BP1 HEK293 cells. Scale bar = 10 µM. (**d**) Stress granule dissolution kinetics after hyperosmotic stress in HEK293 Flp-In T-REx cells at 1 min (left) or 30 min after recovery. Scale bar = 10 µM. (**e**) Schematics of biotin labeling performed in stress BioID timecourse experiments. (**f**) Western blot analysis of puromycin incorporation for 10 min during the indicated stress and recovery periods. NS = No stress, As(III) (sodium arsenite, 500µM, 35 min), NaCl (0.2M, 35 min). (**g**) Overview of stress BioID experiments performed.

**Figure S3. Hyperosmotic stress induces proximal associations with microtubule-associated proteins.** (**a**) Dot plot visualization of selected high-confidence preys identified in DDX3X stress BioID timecourse. As(III) (sodium arsenite, 500µM, 5’-35’), NaCl (0.2M, 45’-75’). (**b**) Schematics show alternative hyperosmotic stress timecourse set up (top). Dot plot visualization of selected high-confidence preys identified by DDX3X during alternative hyperosmotic stress conditions (bottom). (**c**) Heatmap visualization of hierarchical clustering of high-confidence preys identified in DCP1A and DDX3X stress BioID. Baits and conditions are clustered on the x-axis, and high-confidence interactors are clustered on the y-axis based on their abundance.

**Figure S4. DIA analysis improves data quality.** (**a**) Illustration of Data-Dependent Acquisition (DDA) *vs.* Data-Independent Acquisition (DIA) methods in mass spectrometry. (**b**) Diagram of BioID DIA analysis workflow. (**c**) Bar graph of identified proteins in each BioID time-course experiment using DDA and DIA methods. Error bars show variation between replicates. (**d**) Violin plots show the pairwise Jaccard indices between BioID samples. Each point represents a pairwise sample comparison. Statistical significance for three groups of pairwise comparison was evaluated using Mann Whitney U-tests.

**Figure S5. Pearson correlation analysis of processed DIA data.** Heatmap displays the Pearson correlation coefficient calculated between all DIA samples acquired prior to statistical analysis using SAINTq.

**Figure S6. Total proteome changes upon stress, stress-induced proximal interaction changes and validation of interactors.** (**a**) Volcano plot shows total proteome abundance changes in oxidative stress (As (III), sodium arsenite, 0.5mM) or hyperosmotic stress (0.2M NaCl) conditions. Vertical lines indicate 1.5-fold changes. (**b**) The violin plot shows the distributions of abundance fold-changes in high-confidence interactors during stress time course. (**c**) Colocalization validation of identified high-confidence interactors to stress granules in HeLa Flp-In T-REx cells. TRIM56 (endogenous); LARP6-mCherry (stable expression). Scale bar = 10 µM.

**Figure S7. Localization of large ribosomal subunits during oxidative and hyperosmotic stress**. (**a**) Left: Representative images of HeLa cells treated with As(III) (0.5mM sodium arsenite) or 0.2M NaCl for 30 min were permeabilized with 0.01% digitonin, then fixed and processed for immunofluorescence microscopy of RPL32 and G3BP1. Middle: Colocalization is assessed using a line graph on selected stress granules. Right: Images of selected stress granules. Scale bar = 2 µM. (**b**) Left: Representative images of immunofluorescence microscopy of RPL36 and G3BP1, same conditions as panel a. Middle: Colocalization is assessed using a line graph on selected stress granules. Right: Images of selected stress granules. Scale bar = 2 µM. (**c**) Pearson correlation coefficient (PCC) analysis between the RPL32 and G3BP1 channels. A t-test was used to determine the statistical significance of changes in the extent of colocalization between untreated (UT), As(III) and NaCl groups with Holm adjustment (ns, non-significant, n= 2 biological replicates). (**d**) Pearson correlation coefficient (PCC) analysis between the RPL36 and G3BP1 channels. A t-test was used to determine the statistical significance of changes in the extent of colocalization between untreated (UT), As(III) and NaCl groups with Holm adjustment (ns, non-significant, * padj <0.05, n= 2 biological replicates).

**Figure S8. Dynamic Interactions of the CCR4-NOT complex, Protein quantification of CCR4-NOT complex upon knockout or knockdown and the effect of cordycepin on PABPC1. (a)** Volcano plot shows abundance changes in G3BP1 high-confidence interactors during steady to stress state transition. (**b**) CCR4-NOT complex abundance in Cas12a-mediated knockout and/or esiRNA-mediated knockdown HeLa cells used for stress granule quantifications in Figure 3F. Protein abundance information was obtained using DIA mass spectrometry analysis of processed cell lysates collected in no stress condition. Protein abundance is normalized to their abundance in control (no gRNA and scrambled esiRNA) (**c**) Representative images show the effect of cordycepin (50 µM, 2 h) on G3BP1 and PABPC1 during oxidative stress (sodium arsenite (III); 60 µM, 1h) in HeLaFlp-In T-REx cells. Scale bar = 50 µM. (**d**) Violin-Box plot of PABPC1 puncta numbers in HeLa Flp-In T-REx cells pretreated with increasing concentrations of cordycepin for 2 h, followed by no treatment or combined with sodium arsenite (III) stress (60 µM or 75 µM, 1 h). Quantifications from three biological replicates were assessed with Mann Whitney test for statistical significance compared to 0 cordycepin treatment (**** p_adj_ < 10^-4^).

**Figure S9. Subcellular localization of CCR4-NOT complex.** (**a**) Top two rows: Immunofluorescence microscopy of HEK293 Flp-In T-REx cells in no stress or stressed conditions (500 μM sodium arsenite, As(III), 30 min or 0.2 M sodium chloride, 1 h). Bottom panel shows images of HEK293 Flp-In T-REx cells inducibly and stably expressing DCP1A-3xFLAG construct. CNOT1 (green), P-body marker EDC3 (red), nuclei (DAPI). Scale bar = 10 µM. (**b**) Immunofluorescence microscopy of HEK293 Flp-In T-REx cells in no stress or stressed conditions (500 μM sodium arsenite, As(III), 30 min or 0.2 M sodium chloride, 1 h). CNOT1, CNOT7, or CNOT8 (red) paired with stress granule marker G3BP1 (green), nuclei (DAPI). Intensity line plot made in Fiji. Scale bar = 10 µM. (**c**) Immunofluorescence microscopy of CNOT2-3xFLAG stable cells, untreated or stressed for 30 min, stained for stress granule marker (G3BP1) and FLAG tag on CNOT2-3xFLAG construct. Scale bar = 10 µM. (**d**) Live cell images of HeLa stable cells inducibly expressing mNeon tagged CCR4-NOT component and mScarlet-G3BP1, stressed with 500 μM sodium arsenite, As(III) for 30 min. Scale bar = 10 µM.

**Figure S10. CNOT1 sensitivity to cycloheximide.** (**a**) Top: Experimental workflow showing how cells were stressed with 60µM sodium arsenite (As(III)) for 1 h, then treated with carrier or cyclohexamide (CHX, 10μg/mL) for 0.5 h or 1 h. Bottom: Representative immunofluorescence microscopy images show dissolution of stress granules (G3BP1) and CNOT1 upon CHX treatment. Scale bar = 50 µM.

**Figure S11: Global transcriptomic analysis of stress-induced poly(A) lengths and abundance changes dependent on stress granules**. (a) Immunofluorescence microscopy images of HeLa cells in no stress, stressed (sodium arsenate, As(V) 20mM, 1 h) or stress with lipoamide pretreatment (200µM, 1 h pretreatment, then continued for 1 h during stress) conditions stained for endogenous CNOT1 and stress granule marker G3BP1 (green). Scale bar = 50 µM, and 20 µM for enlarged view. (b) Transcriptome-wide distribution of gene-level average poly(A) lengths during no stress (n=3), stress (n=3), lipoamide+stress (n=3), and lipoamide (n=1) with the average values indicated in the middle of each violin plot. Poly(A) lengths were compared for the whole dataset. Mann-Whitney-U test on average of three biological replicates for no stress, arsenate As(V), lipoamide + arsenate As(V), and one replicate for lipoamide alone, Bonferroni correction (**** padj <10^-4^). (c) Density plots showing poly(A) length distributions for JUN and FOS transcripts during no stress, stress (As(V)), and lipoamide + stress (As(V)). TAILcaller is used to determine transcripts with significant changes in poly(A) tail distribution. For the JUN and FOS transcript examples, the Kruskal-Wallis test was employed to determine statistical significance. (d) (Left) Volcano plot of poly(A) length fold-changes from no stress to lipoamide + stress (As(V)) and associated adjusted p-values as determined by TAILCaller and subsequent post-hoc tests. A 1.3-fold cut-off for fold-change was used (n=3). (Right) Volcano plot of abundance fold-changes from no stress to lipoamide + stress (As(V)), and associated adjusted p-values, as determined by DESeq2. A 1.5-fold cut-off for fold-change was used (n=3).

**Figure S12: Analysis of transcripts regulated by CCR4-NOT sequestration to stress granules, and validation of DESeq2**. (a) Dotplot visualization of GO analysis of genes with significant changes during stress, reversed by lipoamide, in both their poly(A) length and abundance (shared changes), poly(A) changes only, or abundance only. Selected GO terms were ordered based on the fraction of genes that belonged to the indicated GO term amongst the total significant list. (b) Bar plot of median poly(A) lengths for 15 genes with significant poly(A) length and abundance changes, which were reversed by lipoamide (n=3, median value taken from all 3 replicates). All genes were significant and made the 1.3-fold-change cut-off in no stress *vs.* stress and stress *vs.* lipoamide+stress, except GAPDH and GADD45A, which show non-significant changes. (c) Bar plot of log_10_(RPM) for 14 genes with significant poly(A) length and abundance changes, which were reversed by lipoamide; two control genes, GAPDH and GADD45A, used for RT-qPCR validation were also included (n=3). All genes were significant and made the 1.5-fold-change cut-off, except GAPDH and GADD45A, which are included as a comparison. (d) RT-qPCR quantification of transcript abundance, testing candidate genes which were significantly increased during stress (As(V)) and reversed by lipoamide (n=3). Abundance is normalized to the ‘unstressed’ condition. Pairwise t-test was performed on three biological replicates, Bonferroni correction (* padj <0.05, ** padj <10^-2^). (e) RT-qPCR quantification of transcript abundance, testing candidate genes which were significantly decreased during stress (As(V)) (n=3). Abundance is normalized to ‘unstressed’ condition Pairwise t-test was performed on three biological replicates, Bonferroni correction (* padj <0.05, ** padj <10^-2^).

**Table S1.** Stress BioID of G3BP1 and FMR1 during oxidative stress and Stress BioID of G3BP1 with lipoamide pre-treatment, related to Figure 1B-C.

**Table S2.** High Information Bait Selection, related to Figure 1D.

**Table S3.** Stress BioID of Nine Stress Granule or Stress Granule& P-body baits during stress and alternative hyperosmotic stress conditions analyzed in DDA mode.

**Table S4.** Stress BioID of Nine Stress Granule or Stress Granule& P-body baits during stress analyzed in DIA mode, related to Figure 2.

**Table S5.** Analysis of Conserved High-Confidence Proximal Interactors, related to Figures 2D-E.

**Table S6.** Linear Mixed Model ANOVA Analysis to Determine Differentially Interacting Proximal Interactors, related to Figure 2F.

**Table S7.** Total Proteome Abundance Changes during Oxidative Stress (0.5mM sodium arsenite, As(III)) and Hyperosmotic Stress (0.2M NaCl) using mixed model ANOVA Analysis to Determine Differential Abundant Proteins, related to Figures S6A.

**Table S8.** Nanopore DRS results, measuring transcriptome abundance and poly(A) lengths, related to Figure 6.

**Table S9.** RT-qPCR primer information, related to Figure S11D-E.

## Notes

### Summary of Updates

Figure 6 and the supplemental files (Supplementary Figures 5, 6, 7, 8, 10, 11, and 12) have been added; the co-authors have been updated.

