## Supplementary figures and images for "Stress Granule Sequestration of CCR4–NOT Promotes Poly(A) Lengthening of Stress-Survival Transcripts"

### Supplemental Figure 1

#### Supplementary Figure 1

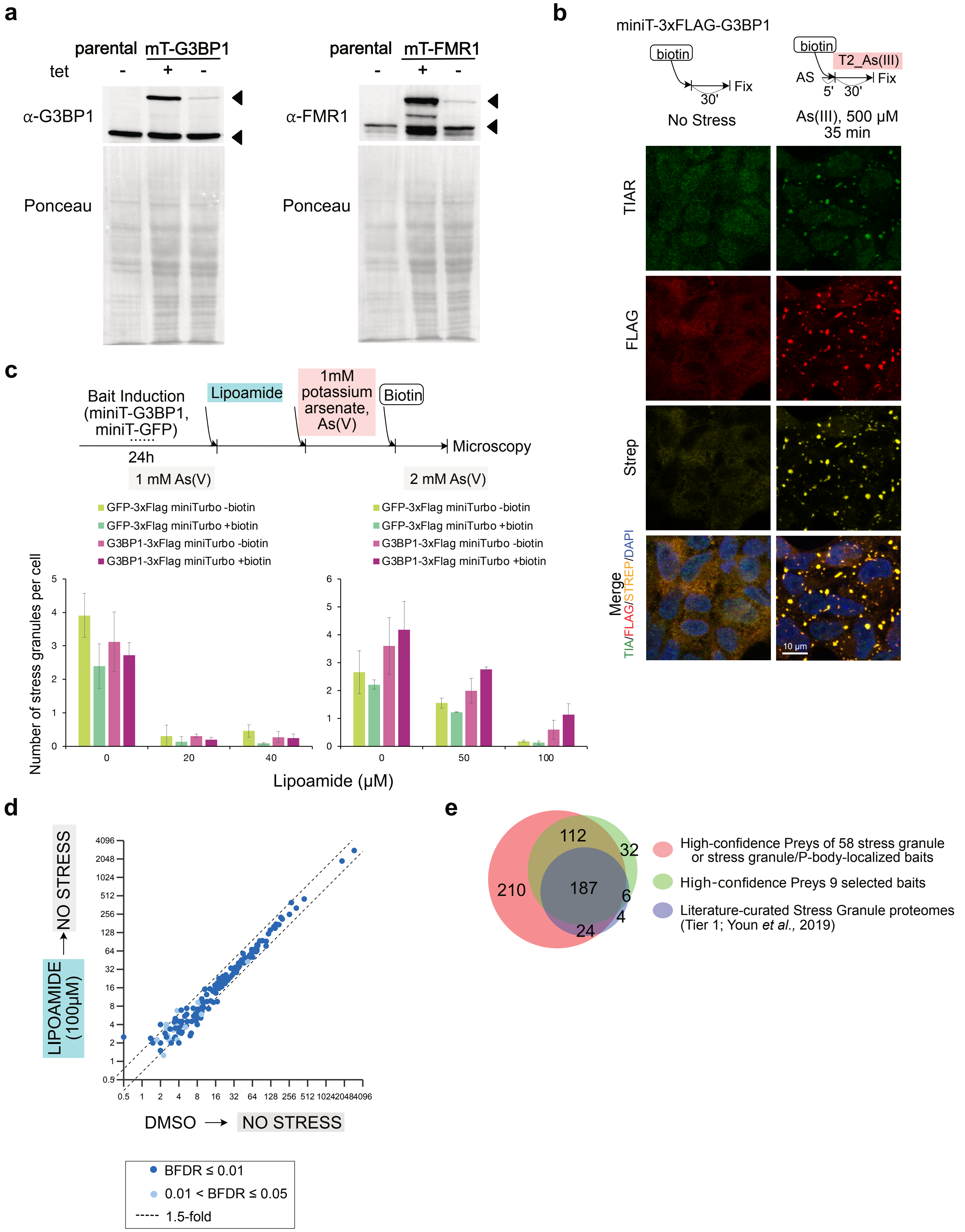

### Supplemental Figure 2

##### Supplementary Figure 2

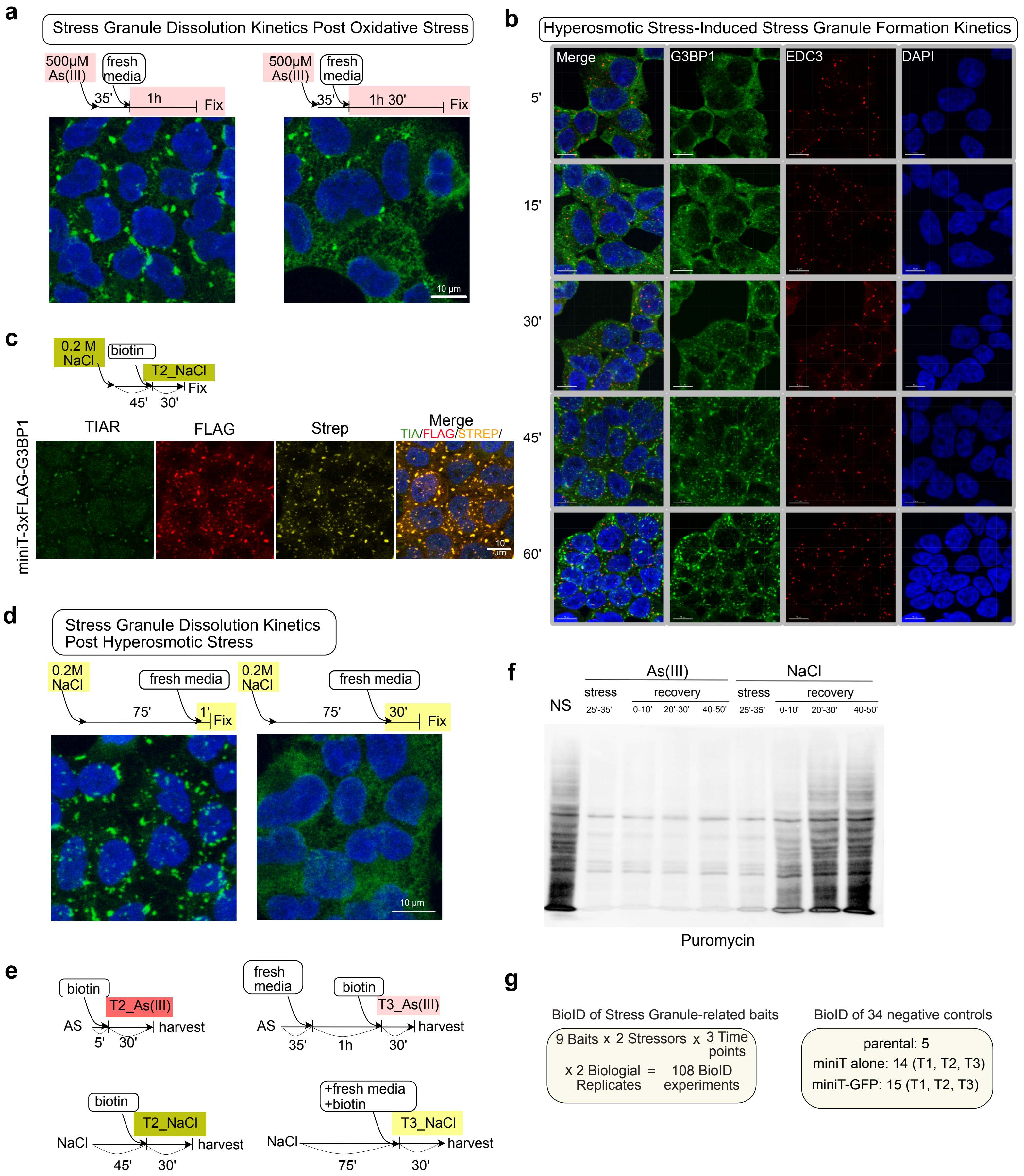

### Supplemental Figure 3

#### Supplementary Figure 3

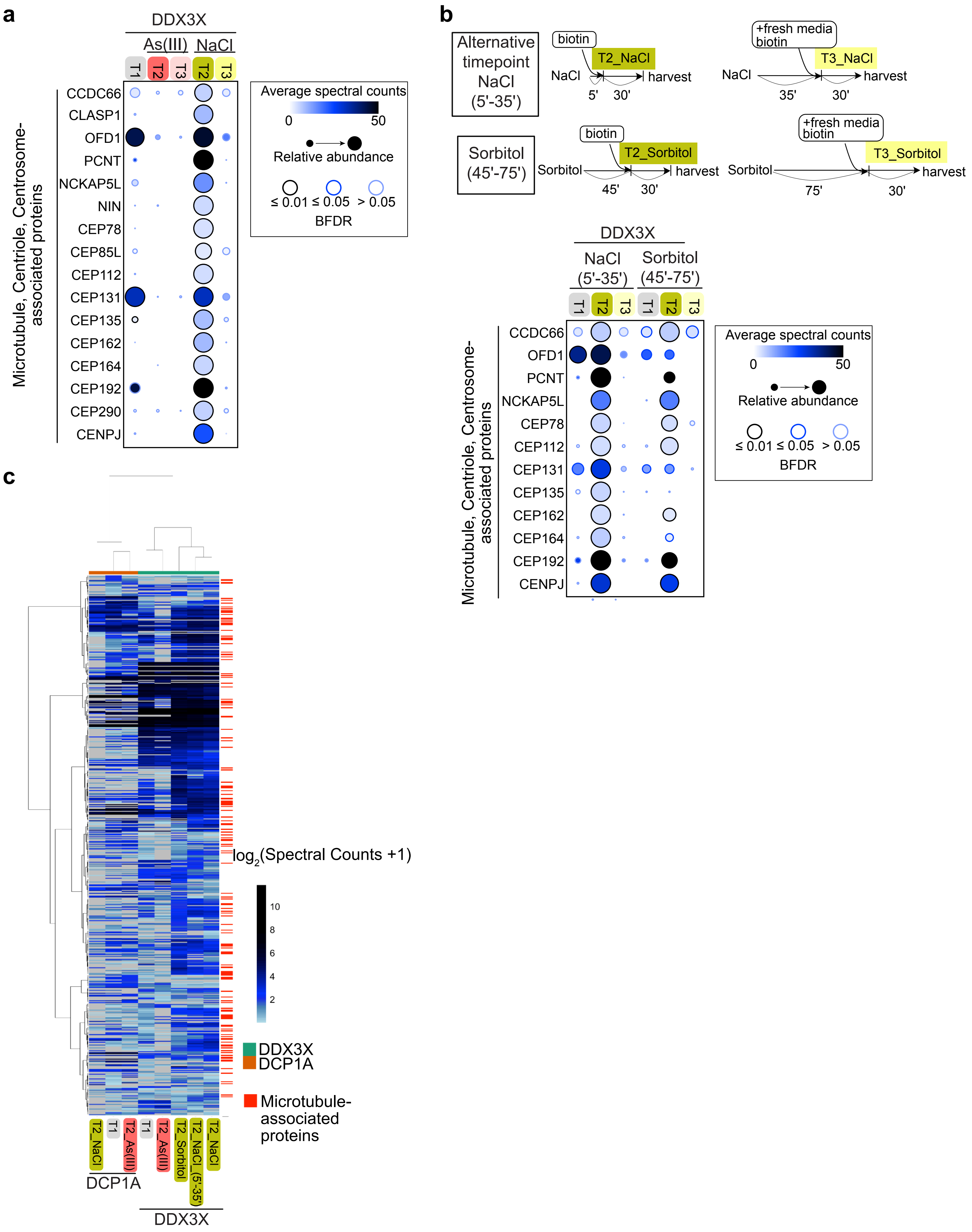

### Supplemental Figure 4

#### Supplementary Figure 4

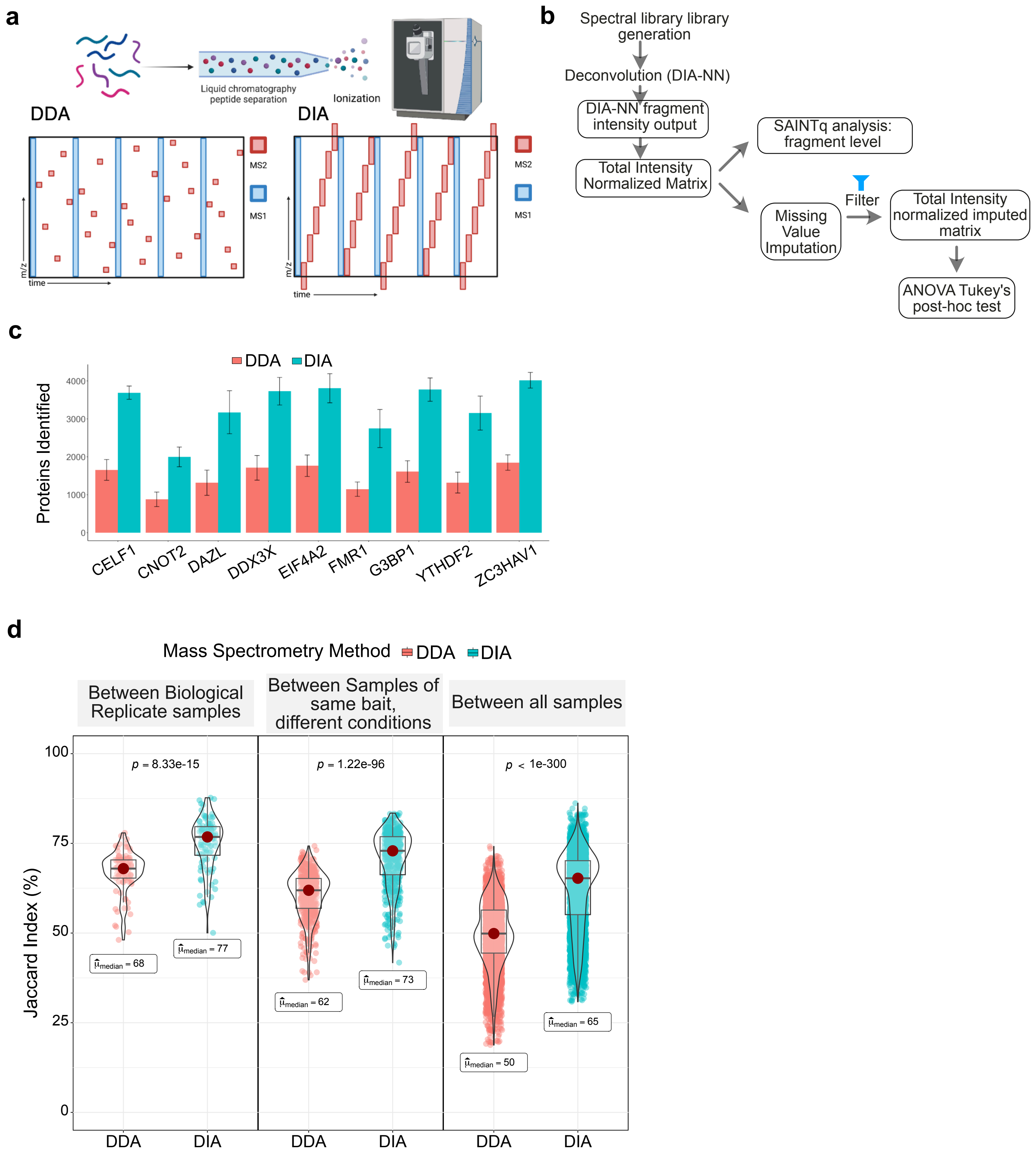

### Supplemental Figure 5

#### Supplementary Figure 5

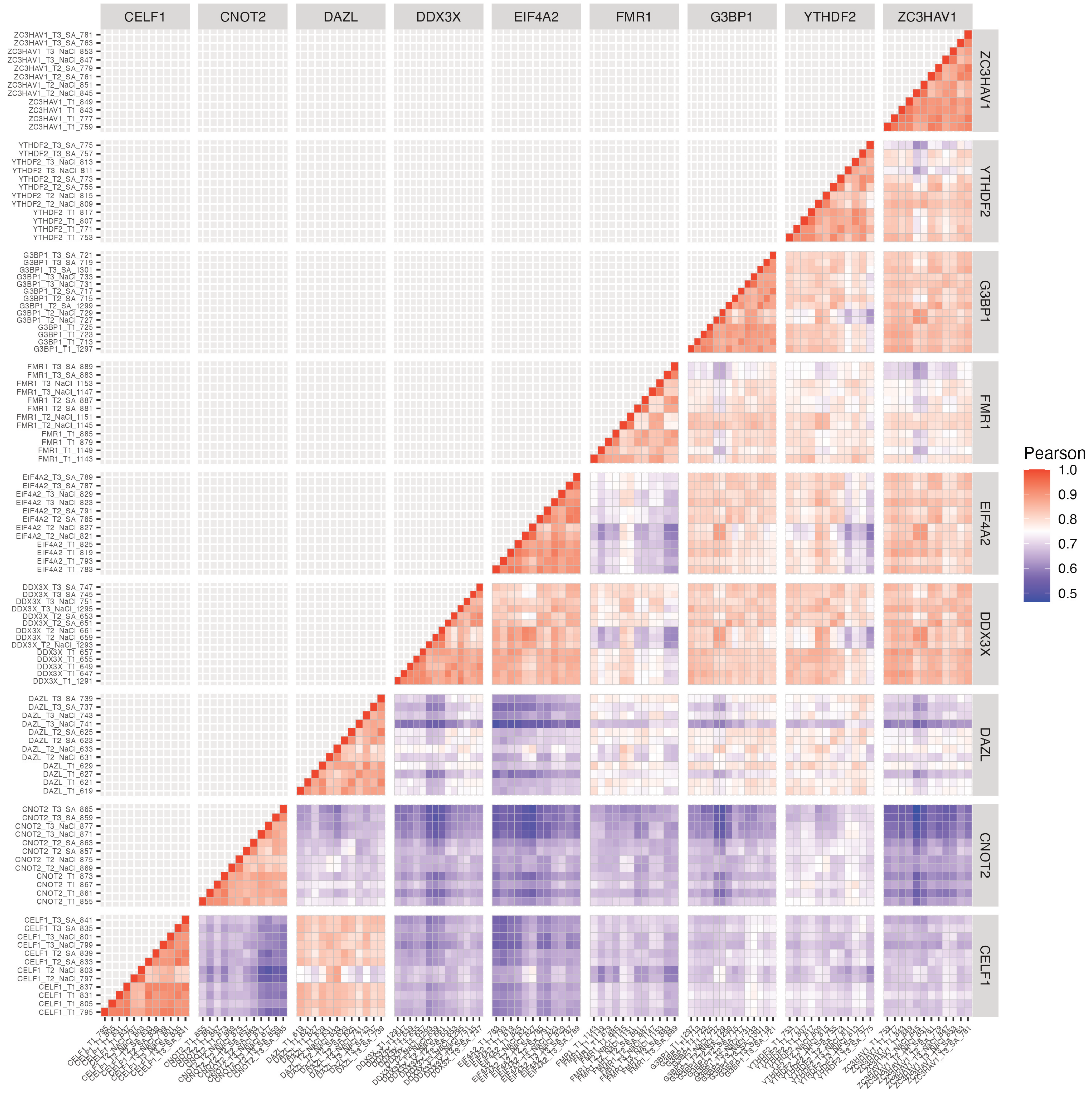

### Supplemental Figure 6

#### Supplementary Figure 6

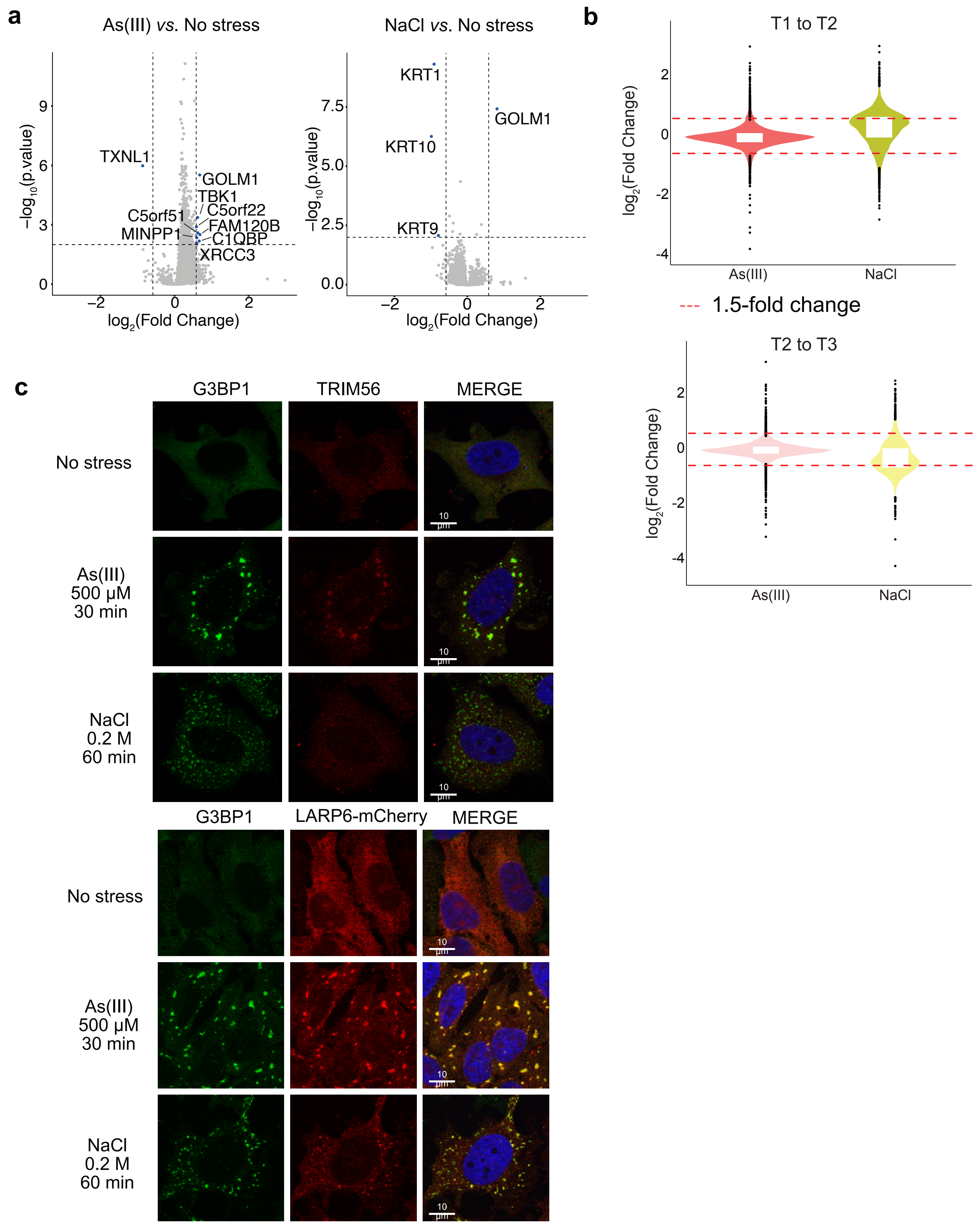

### Supplemental Figure 7

##### Supplementary Figure 7

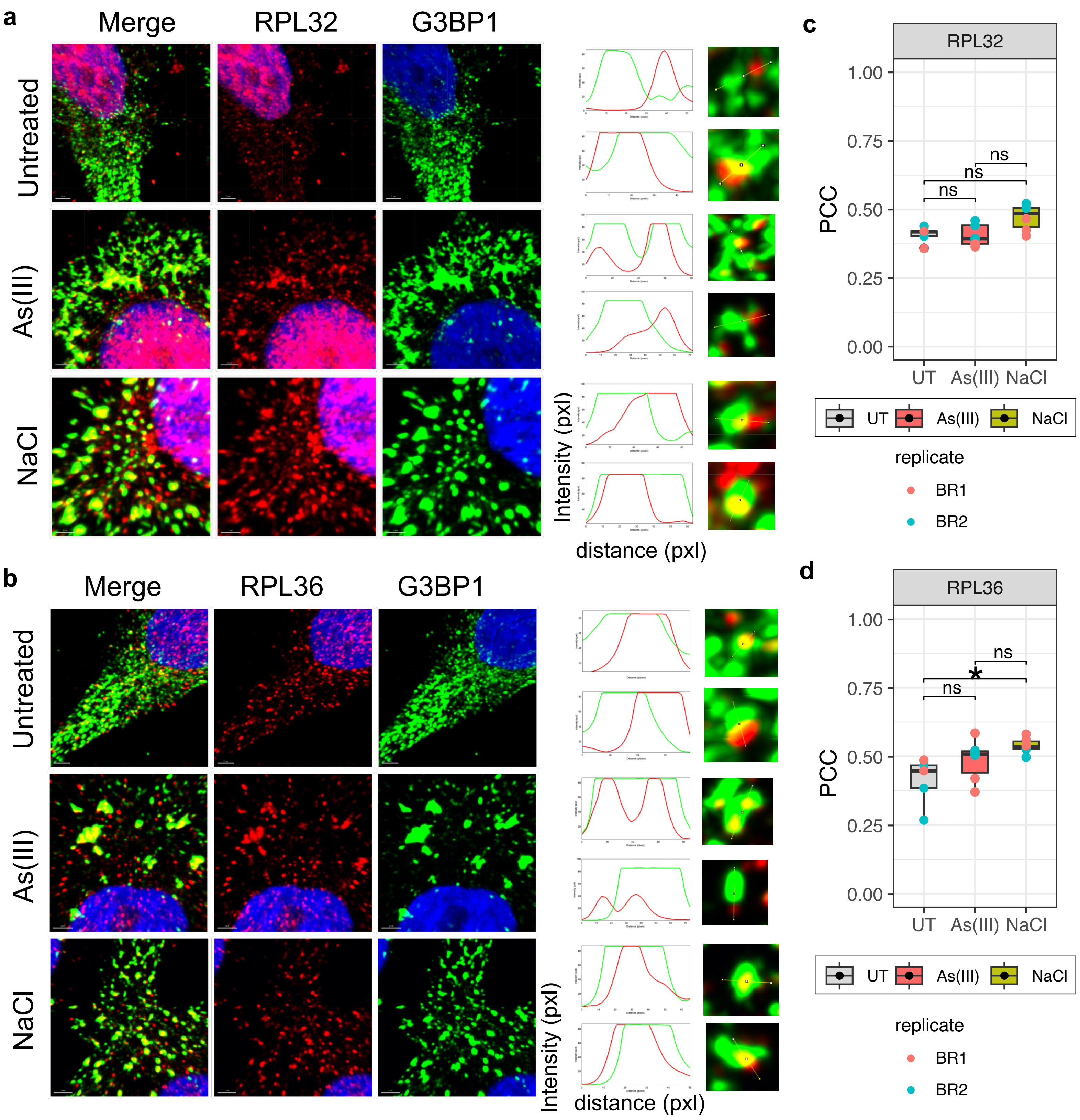

### Supplemental Figure 8

##### Supplementary Figure 8

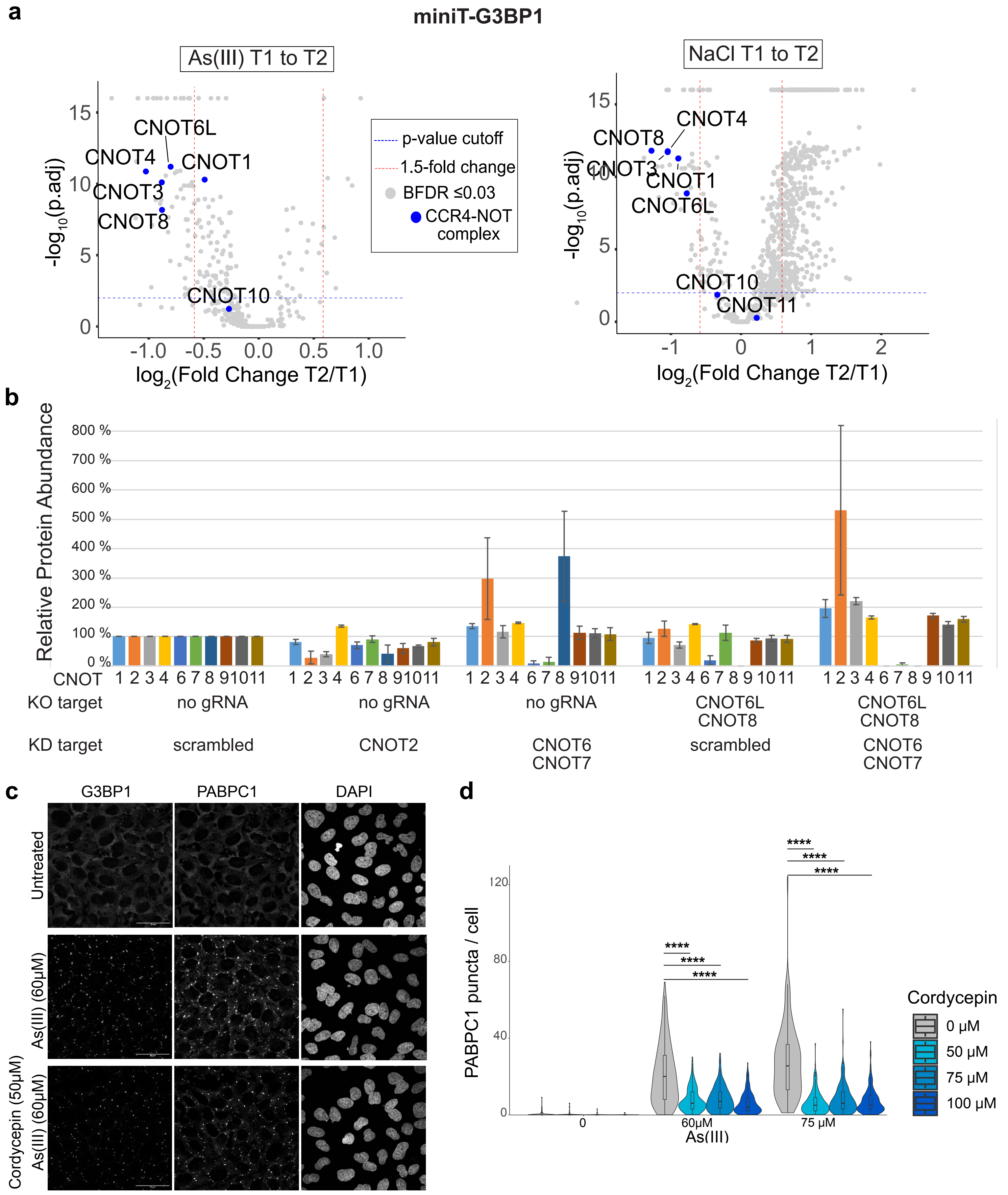

### Supplemental Figure 9

#### Supplementary Figure 9

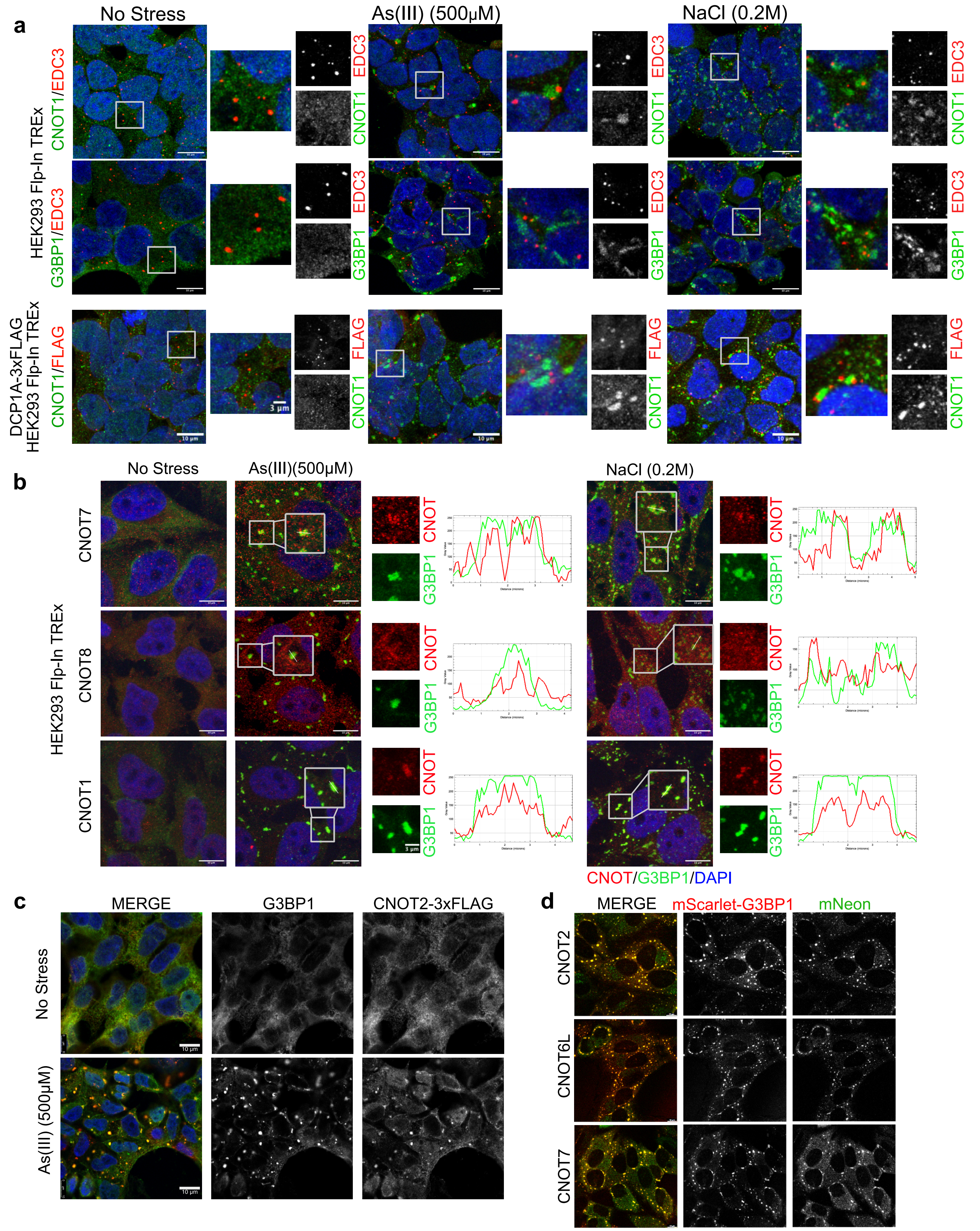

### Supplemental Figure 10

**Supplementary Figure 10**

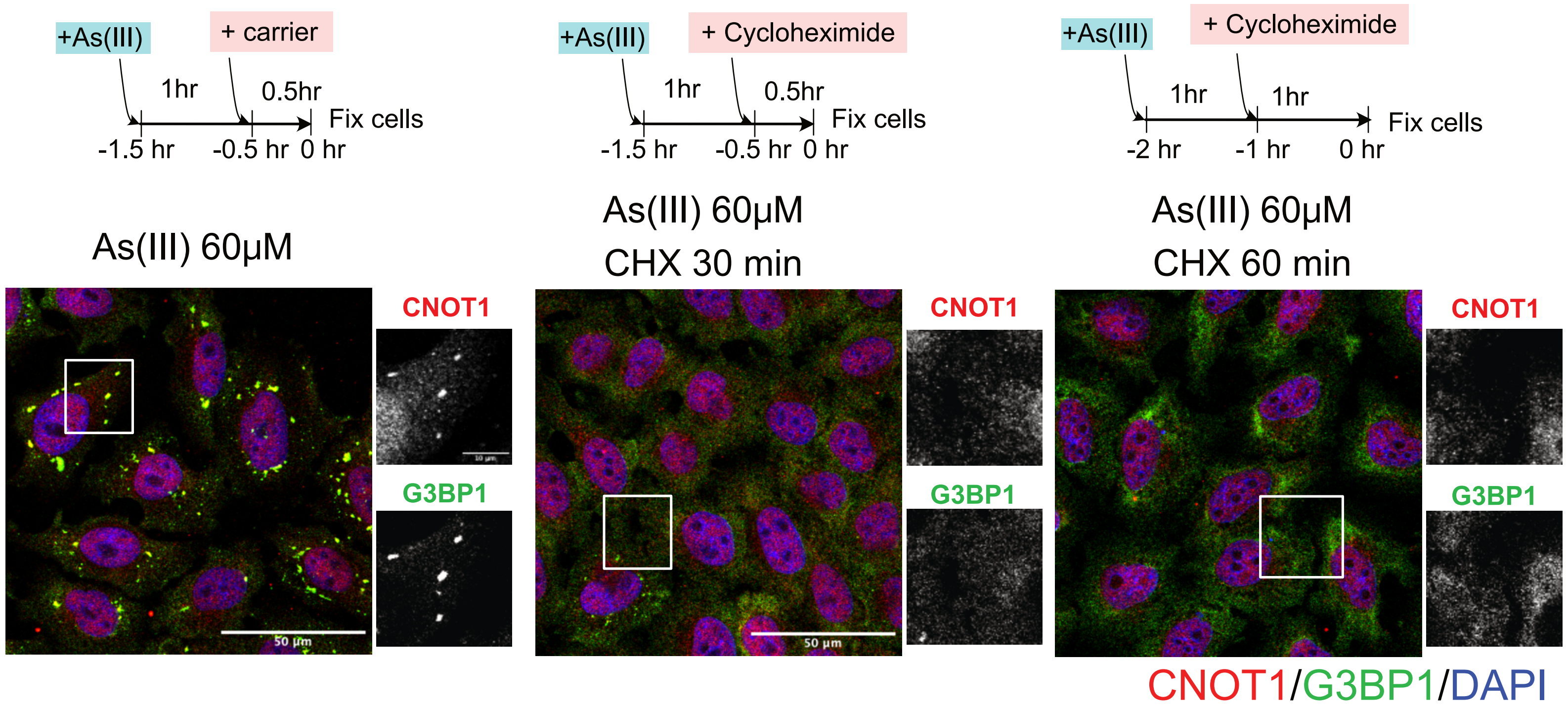

### Supplemental Figure 11

#### Supplementary Figure 11

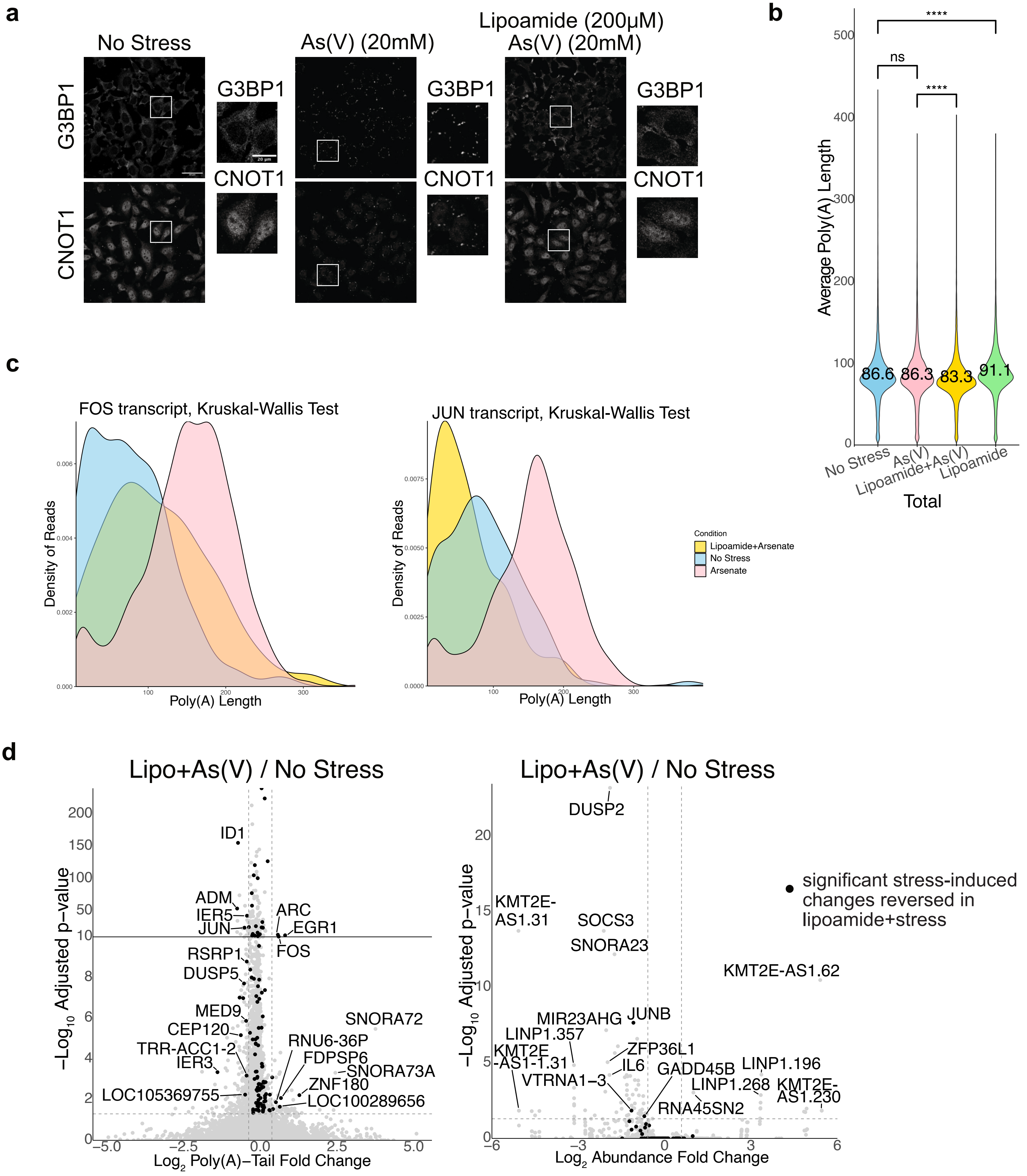

### Supplemental Figure 12

#### Supplementary Figure 12

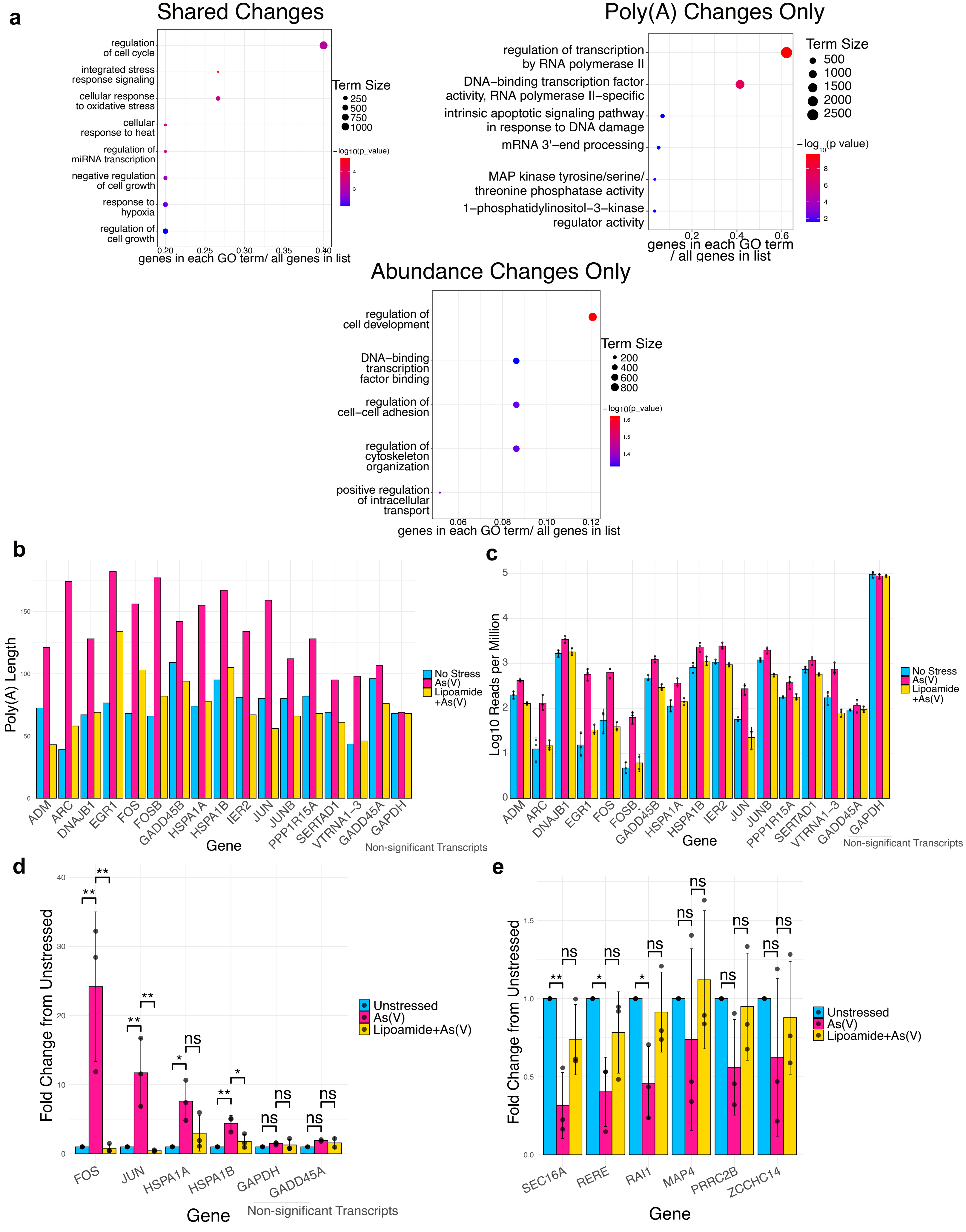
